# Unlocking meiotic crossovers near plant centromeres

**DOI:** 10.1101/2025.08.19.671075

**Authors:** Rigel Salinas Gamboa, Joiselle Blanche Fernandes, Qichao Lian, Dipesh Kumar Singh, Stephanie Durand, Divyanshu Sahu, Laia Capilla Perez, Hernan Lopez, Alexandra Kalde, Raphael Mercier

**Affiliations:** Department of Chromosome Biology, Max Planck Institute for Plant Breeding Research, Carl-von-Linné-Weg 10, 50829 Cologne, Germany; Guangdong Laboratory for Lingnan Modern Agriculture, Guangdong Provincial Key Laboratory for the Development Biology and Environmental Adaptation of Agricultural Organisms, College of Life Sciences, South China Agricultural University, Guangzhou 510642, China; Laboratory of Plant Reproduction, Center for DNA Fingerprinting and Diagnostics (CDFD), Uppal, Hyderabad 500039, India

## Abstract

Crossovers (COs) ensure proper chromosome segregation during meiosis and generate genetic diversity. COs are non-uniformly distributed along chromosomes and almost universally suppressed in centromere-proximal regions, notably creating a significant bottleneck for plant breeding. The mechanism of this CO suppression is elusive, but chromatin state is a contributing factor. Here, we identify three factors that actively limit proximal CO in Arabidopsis, the cohesion establishment factor CTF18, the centromeric cohesin protector SGO2, and the deSUMOylase SPF2. The mutation of these factors allows both the formation of CO in centromere-proximal region where they were completely absent in the wild-type, and enhance their frequency where they were rare. COs can be further increased by combining these mutations together or with mutation in the DNA methylase *CMT3*, suggesting that multiple mechanism prevent proximal CO in parallel. The identification of the very conserved CTF18, SGO2 and SPF2 as suppressors of centromere-proximal CO highlights the importance of cohesin turnover in this process and opens up new possibilities for plant breeding.

## Introduction

Centromeres are specific regions of chromosomes where the kinetochore complex attaches, enabling microtubule binding and chromosome segregation during mitosis and meiosis ^1^. In many eukaryotes, centromeres contain the histone variant CENH3/CENPA and can be small as a single nucleosome as in budding yeast, a few kilobases (Kb) long as in fission yeast, or constituted of several megabases (Mb) of satellite arrays like in Human (171bp alpha-satellite) and Arabidopsis (CEN178 satellite) ^2,3^. In many species, centromeres are surrounded by regions with low gene density, high transposon density, rich in DNA methylation and heterochromatin marks, and suppressed meiotic recombination. These centromere-proximal regions are commonly referred to as pericentromeres. However, there is no consensus in the literature on defining the boundaries of pericentromeres, either considering gene density, DNA/histone methylation levels, or reduced recombination rates ^3–8^.

Meiotic crossovers (COs) enhance genetic diversity and are also essential for accurate chromosome segregation at meiosis. By creating exchanges between homologous chromosomes, COs physically link them and promote their balanced segregation. Meiotic COs results from the repair of programmed DNA double-strand breaks (DSBs). Only a small subset of DSBs are repaired as homologous COs; the rest is repaired using either the sister chromatid as a template, or the homologous chromatid but resulting in non-CO events ^9–12^. The distribution of COs along chromosomes is non-uniform, with regions prone to CO and other regions where they are rare or absent, most notably in the pericentromeric regions ^13–17^. CO suppression at the centromere-proximal region, the so-called “centromere effect,” was initially described in *Drosophila* ^18^ and is conserved across species, from unicellular to multicellular eukaryotes, including yeast, plants and animals ^19,20^. CO suppression applies to the centromere itself (e.g. the satellite array) and extends to the flanking pericentromeres at varying distances between species. In budding yeast with a point centromere, CO suppression ranges in different chromosomes from ∼1 kb to 18 kb ^21^. In rice, Arabidopsis, Drosophila and humans, the region lacking COs extends from a few 100kbs to a few Mbs beyond the centromere ^16,22–26^. In many crop species, the pericentromeric COs suppression extends to more than half of the chromosomes, representing tens of Mbs of DNA in tomato or maize, and a hundred Mbs in wheat ^5,27–30^.

This centromere-proximal CO suppression is thought to be beneficial, as COs near centromeres have been associated with an increased risk of chromosome mis-segregation in humans, Drosophila, and yeast ^2,31–34^. It is suggested that CO and thus chiasmata formation closer to the centromere would locally weaken sister chromatid cohesion, which is essential for balanced sister chromatid distribution at second meiotic division. Alternatively, proximal COs could generate an entanglement preventing chromosome separation at anaphase I. However, these hypotheses cannot easily explain the extension of CO suppression to large proportions of the chromosomes.

Increasing centromere-proximal COs could positively impact plant breeding. As described above, in many crops, large regions spanning megabases of DNA near centromeres are devoid of COs ^5,35–38^. Even if gene density tends to be reduced, the centromere-proximal region contains genes that are thus inaccessible to breeders. For example, 18% of the barley genes are located in the low-recombining centromere-proximal region ^4^. The wheat pericentromeric region contains crucial defense response genes ^5,39^. The Tomato Mosaic Virus (ToMV) resistance gene (*Tm-2*^2^) was introgressed from a wild species into commercial varieties, together with a 64Mb non-recombining pericentromeric chromosomal segment that alters multiple tomato traits ^40–43^. In potatoes, inbreeding depression causes harmful mutations to accumulate near centromeres due to low recombination rates ^44^. Low COs in centromere-proximal regions thus challenge breeding by limiting the combination of favorable alleles and counterselection of harmful ones.

Although the centromere effect was described almost a century ago ^18^, the mechanisms underlying this CO suppression remain unclear. Pericentromeres are enriched in heterochromatin marks, including DNA methylation, lysine 9 of histone H3 methylation (H3K9me2), and histone variant H2A.W, which contribute to CO suppression. Supporting this conclusion, mutations in the H3K9 methyltransferase genes in *Drosophila* (*Su(var)3-9*) and fission yeast (*clr4*) increase centromere-proximal DSBs and COs. In Arabidopsis, disruption of the H3K9 methyltransferase KYP/SUVH4 SUVH5 SUVH6, or the non-CG DNA methyltransferase CMT3, or the histone variant H2A.W also leads to an increase in pericentromeric COs ^15,45–48^. However, the absence of the CG methylation factor MET1 in Arabidopsis has the opposite effect, a decrease in centromere-proximal COs ^15,49^. This contrasting effect may be due to the differences in the heterochromatin mark H3K9me2, which is reduced in *cmt3* mutants but remains intact in *met1* mutants ^50,51^. Besides chromatin factors, kinetochores and cohesins have been implicated in CO suppression in yeast. In budding yeast, disruption of subunits of the inner kinetochore complex Ctf19/CCAN increases both COs and NCOs in pericentromeres ^52^. Ctf19 plays a dual role in suppressing centromere-proximal COs: (i) CTF19 limits DSB formation at the centromere-proximal region. (ii) CTF19 recruits and enriches the REC8-cohesin complex, which in turn prevents the repair of DSBs as COs ^52–54^. In fission yeast, DSB formation depends on the Rec8-Rec11 cohesin complex, which tends to be replaced by the Rec8-Psc3 complex in the pericentromeric region, preventing DSBs and thus CO formation ^2,55^. This enrichment of Rec8-Psc3 in the centromere-proximal region is dependent on the H3K9me-binding protein Swi6 ^2^. Finally, the transverse element of the synaptonemal complex (which zips the homologs together during meiotic prophase) Zip1, was shown to contribute to pericentromeric CO suppression in budding yeast ^52^.

In this study, we performed a genetic screen in *Arabidopsis thaliana* and identified three factors preventing centromere-proximal COs: the cohesion establishment factor CTF18, the centromeric cohesion protection factor Shugoshin (SGO2), and the deSUMOylase SPF2. In each corresponding mutant, we observed an increase in pericentromeric COs, including regions where COs are absent in the wild-type. This establishes the role of cohesin regulators in preventing COs near centromeres in plants. We showed that the induced centromere-proximal COs occur in gene islands present in syntenic regions. We also show that proximal COs can be further increased by combinations of these mutations and disruption of *CMT3*, suggesting that multiple mechanisms contribute to CO suppression in parallel. Importantly, unlocking centromere-proximal COs does not significantly affect fertility, making this a promising strategy for accessing genes near centromeres and offering new opportunities for crop improvement.

## Results

### Identification of Mutants with Increased Centromere-Proximal Crossovers

We aim to identify factors that limit COs in the centromere-proximal regions. We selected candidate mutants (see below) and measured recombination rates in intervals flanking centromeres using two methods, looking for mutants with enhanced recombination: (1) We used Fluorescent Tagged Lines (FTL) lines, also known as TRAFFIC lines, containing two genetically linked transgenes that express fluorescent proteins in seeds (GFP and dsRed). Scoring seed fluorescence allows measuring recombination in the interval defined by the position of the two transgenes ^56^ (Figure 1A). We used the line CTL3.437.55, which contains the transgenes CG437 (Col_CEN Chr3:10,942 Kb) and CR55 (Chr3: 18,255 Kb), defining an interval of 7.3 Mb spanning the centromere of chromosome 3, including a 2.1 Mb *CEN178* satellite array ^8^ (Figure 1A). We crossed candidate mutants with CTL3.437.55 and selected plants from the F2 populations that were heterozygous for the fluorescent transgenes and either wild-type or homozygous for the tested mutation. Recombination was measured based on the segregation of fluorescence in seeds from the self-fertilization of mutant and wild-type sister plants. (2) When candidate mutations were available in two different strains, we measured recombination in hybrids (Col/L*er*, Col/Ws) by PCR-genotyping genetic polymorphisms flanking centromeres (Figure S2A-B) in progeny of F1 hybrids, either wild-type or homozygous for the tested mutation.

**Figure 1:**
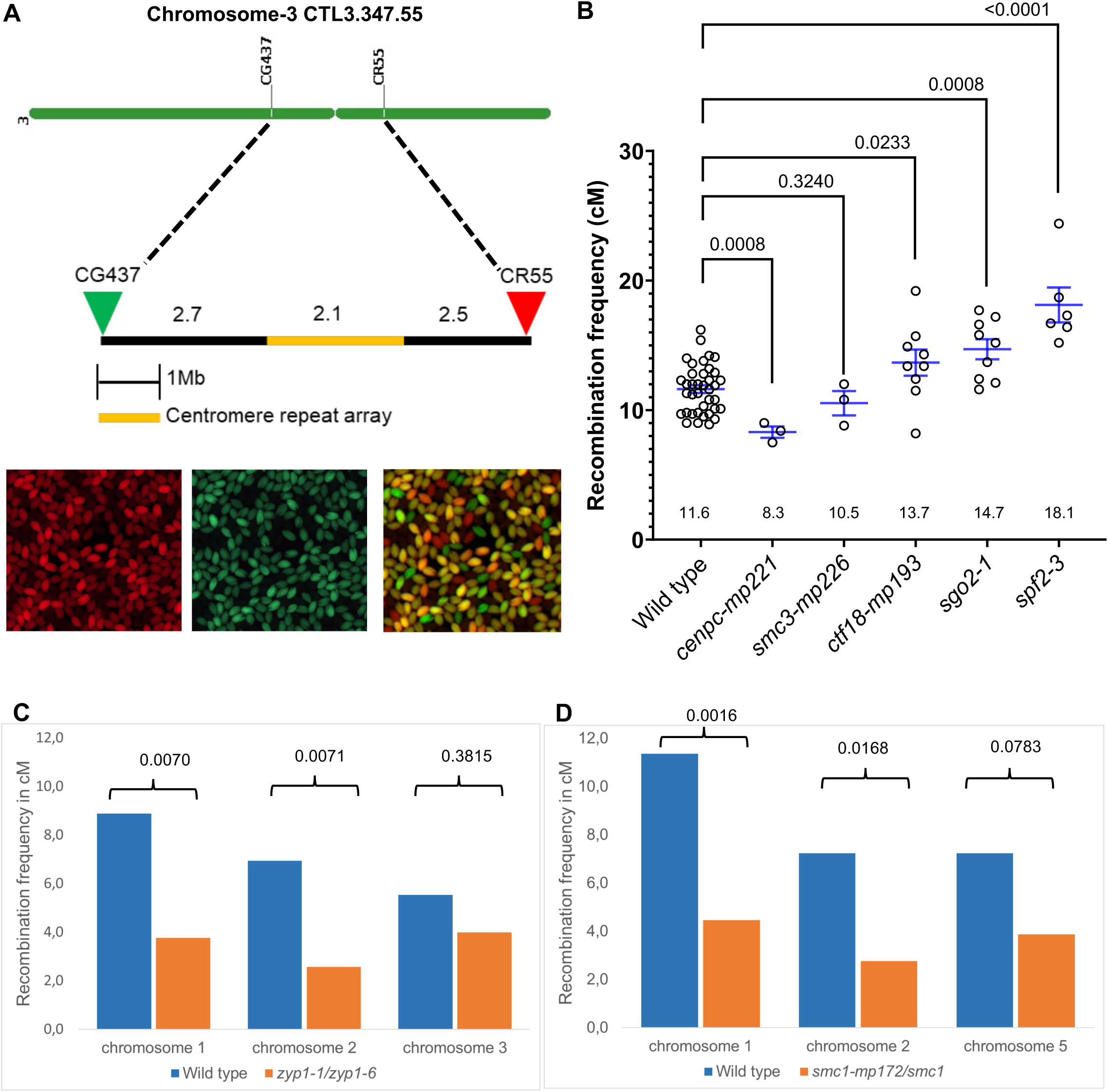
Testing candidate mutations for elevated crossovers in pericentromeres. **(A)** The physical interval on chromosome 3 defined by the transgene conferring green (CG437) and red (CR55) fluorescence to seeds in the line CTL.3.437.55. Physical sizes are indicated in Mb. Seeds showing segregation of the GFP and dsRED fluorescence. (B) Measure of recombination frequency in centiMorgans (cM) in the CTL.3.437.55 interval. Each dot represents a measurement on ∼500 seeds. Blue lines indicate the mean of the recombination frequency +/- standard Error (SEM). Wild-type samples from different populations (i.e., sister plants of different mutants) are pooled together. Measures for individual populations are provided in Figure S2. P-values are from Mann-Whitney tests. (**C**-**D**) Recombination frequencies (cM) measured in 3 centromeric intervals in *zyp1* (Col/Ler) and *smc1* (Col/Ws) hybrids. Schematic representation of centromeric intervals are provided in Figure S2A and S2B. P values are Fisher’s exact tests. Source data for Figures 1A,1B,1C and 1D is provided as supplementary material.

One landmark of meiosis is the orientation of sister kinetochores toward the same pole at meiosis I, which allows the distribution of homologous chromosomes rather than sister chromatids ^57^. Our group identified factors that promote the monopolar orientation of kinetochores at meiosis in Arabidopsis through a forward genetic screening (Singh et al., BioRxiv 2025). This screen uncovered mutant alleles of genes encoding cohesin subunits (SMC1, SCC3, SMC3, REC8), cohesion establishment factors (CTF18, DCC1), protectors of centromeric cohesion (SGO2), an inner kinetochore protein (CENP-C), and a deSUMOylase (SPF2) (Singh et al., BioRxiv 2025) (Figure S1). Importantly, while monopolar orientation and/or cohesion are weakened in these mutants, correct chromosome segregation is maintained. We reasoned that the changes in the organization of the centromere-proximal region that affect kinetochore orientation in the identified mutants may also affect crossover formation near centromeres. Therefore, we tested these candidates for modified recombination in intervals spanning the centromere.

Among the mutants affected in monopolar orientation were *cenpc-mp221*, a leaky allele of CENP-C, an essential kinetochore subunit (Singh et al., BioRxiv 2025)^58,59^. In *cenpc-mp221*, we observed reduced recombination in the tested interval on chromosome 3 when compared to the wild-type (Figure 1B and S2C), contrasting with previous observations in S*. cerevisiae* kinetochore mutants ^52^. Next, we tested two alleles of cohesin subunits, *smc1-mp172* and *smc3-mp226* (Singh et al., BioRxiv 2025). SMC1 and SMC3 are essential subunits of the cohesin complex, which, together with SCC1/REC8 and SCC3, form a ring-shaped structure that holds together sister chromatids at mitosis and meiosis and organizes chromatin folding ^60–62^. Recombination was not significantly modified in *smc3-mp226* in the CTL3.437.55 interval (Figure 1B and S2D). In *smc1-mp172*, two of the three tested intervals showed significantly reduced recombination, while the third interval appeared reduced but not significantly (Figure 1D). This reduced recombination points to a role of cohesins in promoting CO formation, which is also supported by the Arabidopsis *rec8* and *scc3* mutants in which meiotic DSB repair is strongly affected ^63^. Thus, the *cenpc*, *smc1,* and *smc3* alleles tested here do not support a role for the kinetochore or the cohesin subunits in limiting pericentromeric COs.

In yeast and drosophila, the transverse/central element of the synaptonemal complex contributes to the suppression of centromere-proximal COs ^7,52,64^. However, mutation of *ZYP1* – the transverse element in *Arabidopsis* -, leads to a significantly lower CO frequency in two of the three centromeric intervals that we tested (Figure 1C). This is consistent with genome-wide data showing distal redistribution of COs in *zyp1* mutants ^65^ and suggests that ZYP1 does not contribute to suppressing pericentromeric COs but rather favors them.

Next, we investigated loss-of-function mutants of CTF18 and SGO2 which also support monopolar orientation of sister chromatids (Singh et al., BioRxiv 2025). CTF18 is a conserved replication factor that favors the establishment of sister chromatid cohesion ^66–68^. Shugoshins (SGOs) are conserved protectors of centromeric cohesins ^69^. Arabidopsis SGO1 and SGO2 are partially redundant for this function at meiosis ^70,71^. In both *ctf18-mp193* and *sgo2-1,* we observed a ∼20% significant increase in recombination in the CTL3.437.55 interval compared to the wild-type (Figure 1B, S2E-F, Source data). This suggests that the cohesin regulators CTF18 and SGO2 contribute to the suppression of pericentromeric COs.

Finally, we tested another mutant affecting monopolar orientation, *spf2*, a loss-of-function allele of SPF2 (Singh et al., BioRxiv 2025). SPF2 is one of the two homologs of the yeast deSUMOylase Ulp2, which notably regulates cohesin turnover ^72,73^. In *spf2*, recombination was increased by ∼50% compared to wild-type controls in the CTL3.437.55 interval (18.1cM vs 11.6cM) (Figure 1B, Figure S2G). This suggests a pivotal role for the deSUMOylase SPF2 in the restriction of centromere-proximal COs.

In brief, we identified three genes: *CTF18, SGO2*, and *SPF2,* whose loss-of-function mutations led to an increase in CO frequency in the pericentromeric CTL3.437.55 interval, which is particularly marked in *spf2*. This suggests that CTF18, SGO2, and SPF2 negatively regulate CO formation in the pericentromeres, and motivated a comprehensive study of the distribution of CO in their absence.

### Genome-wide analysis confirms the role of *CTF18, SGO2*, and *SPF2* in preventing CO in pericentromeres

To assess the impact of *ctf18, sgo2, and spf2* on COs across the genome, we generated novel mutant alleles in a second strain (Ler-0) using CRISPR-Cas9 (Figure S1). We crossed Col and Ler heterozygous mutants to create *ctf18 -/-*, *spf2-/-*, and *sgo2-/-* Col/Ler F1 hybrids, which were reproduced by self-fertilization. Wild-type +/+ plants were selected in the same F1 populations and also reproduced by selfing. The resulting F2 plants were sequenced by Illumina to detect COs using Col/Ler polymorphisms. To gain power, we combined our 472 wild-type F2 plants with datasets from previous studies using the same Col/L*er* hybrid, making a total of 2,365 F2s (Figure S3) ^15,74^. In addition, a previous study showed that a mutation in *CMT3*, which maintains non-CG DNA methylation, increases centromere-proximal COs ^15^. We reanalyzed the available *cmt3* raw sequence data, which are also Col/Ler F2 samples, with the same parameters and reference genome.

First, we defined the regions where COs are suppressed in the wild type. The Non Recombing Zone (NRZ) was previously defined as a contiguous region flanking the centromere where no COs are observed in the wild type ^8^. As we included more samples than in the previous study, we slightly redefined the NRZ borders (Table 1) (19,167 COs in 2365 F2, equivalent of 4,730 gametes. Source data). This NRZ encompasses the centromeric repeat arrays, which are completely devoid of COs. Note that even if the arrays of centromeric repeats do not contain usable markers, a CO in these regions would be detected because of the genotype switch of the flanking markers (e.g. Col/Col on one side and Col/L*er* on the other side). None of the 19,164 COs identified in 4,730 gametes were mapped to the centromeric repeat regions, suggesting that COs are extremely rare or absent in the *CEN180* repeats arrays that represent a total of 12.6 Mb of DNA, 9.6% of the genome. The NRZ extends beyond the centromere repeats for all five chromosomes, adding 3.7 Mb of CO-free DNA. The NRZ represents a total of 16.3 Mb or 12.3% of the genome (Figure 2, Figure 3, Table 1). Flanking the NRZ, we designated two zones of Low Recombining Zone, LRZ1 and LRZ2, being a physical non-overlapping interval, each corresponding to a genetic size of 1cM in wild type (0.5 cM on each side of the centromere) (Figure 2B). From centromere outwards, LRZ1 immediately flanks the NRZ, and LRZ2 flanks LRZ1. Altogether, the LRZ1 regions of the five chromosomes span 9.8 Mb (and, by definition, 5 cM), with a CO rate of 0.5 cM/Mb, a ∼8-fold reduction compared to the rest of the genome (non-NRZ/LRZ). LRZ2 regions span 2.41 Mb with a CO rate of 2 cM/Mb, a ∼2-fold reduction compared to the rest of the genome (Table 1, Figure 3A).

**Table 1.** Properties of the Non-Recombining Zone (NRZ) and the Low Recombining zones (LRZs).

|  | border position (Mb) |  |  |  |  |  |  |  | Size (Mb) |  |  |  |  | Crossover rate (cM/Mb) |  |  |  |  |
| --- | --- | --- | --- | --- | --- | --- | --- | --- | --- | --- | --- | --- | --- | --- | --- | --- | --- | --- |
|  | *Centrom.*<br>*repeats*<br>*left* | *Centrom.*<br>*repeats*<br>*right* | *NRZ*<br>*Left* | *NRZ*<br>*right* | *LRZ1*<br>*Left* | *LRZ1*<br>*right* | *LRZ2*<br>*Left* | *LRZ2*<br>*right* | *chrom.* | *Centro.*<br>*repeats* | *NRZ* | *LRZ1* | *LRZ2* | *whole*<br>*genome* | *NRZ1* | *LRZ1* | *LRZ2* | *non-*<br>*NRZ/LRZ* |
| Chr1 | 14,84 | 17,56 | 14,19 | 17,72 | 13,47 | 18,24 | 13,29 | 18,26 | 32,5 | 2,72 | 3,53 | 1,24 | 0,20 | 3,04 | 0,00 | 0,81 | 5,02 | 3,52 |
| Chr2 | 3,72 | 5,95 | 3,11 | 6,47 | 2,71 | 7,27 | 2,45 | 7,92 | 22,2 | 2,22 | 3,36 | 1,20 | 0,90 | 3,25 | 0,00 | 0,83 | 1,11 | 4,19 |
| Chr3 | 13,60 | 15,73 | 13,54 | 16,57 | 11,99 | 17,20 | 11,81 | 17,63 | 25,7 | 2,14 | 3,03 | 2,18 | 0,61 | 2,93 | 0,00 | 0,46 | 1,63 | 3,69 |
| Chr4 | 4,20 | 6,98 | 3,92 | 7,16 | 1,64 | 7,85 | 1,54 | 8,12 | 21,6 | 2,77 | 3,24 | 2,97 | 0,37 | 3,07 | 0,00 | 0,34 | 2,70 | 4,29 |
| Chr5 | 11,78 | 14,55 | 11,75 | 14,87 | 10,81 | 16,13 | 10,60 | 16,24 | 29,5 | 2,77 | 3,11 | 2,20 | 0,32 | 3,13 | 0,00 | 0,45 | 3,10 | 3,79 |
| all |  |  |  |  |  |  |  |  | 131,6 | 12,62 | 16,28 | 9,79 | 2,41 | 3,08 | 0,00 | 0,51 | 2,07 | 3,83 |

**Figure 2:**
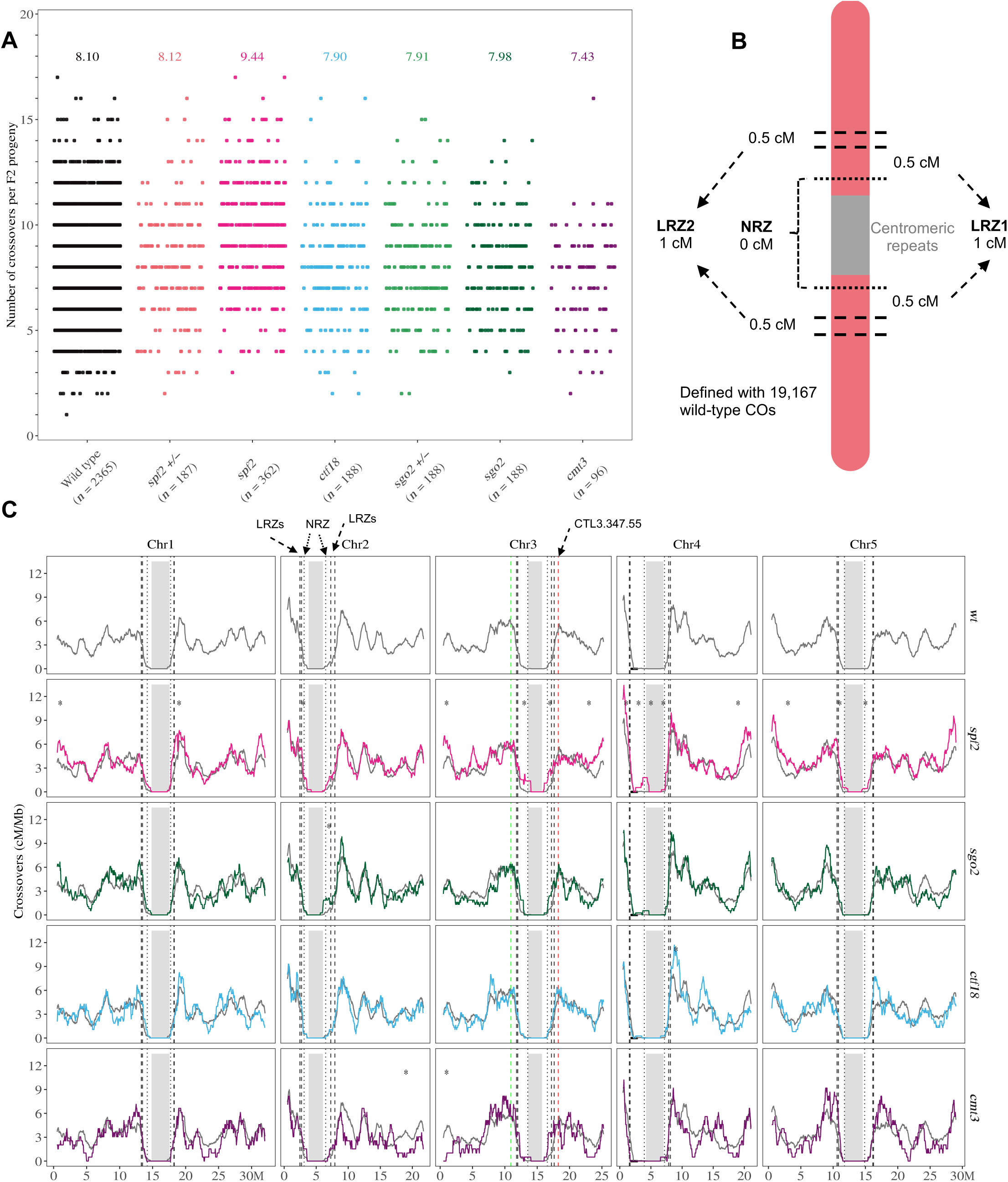
Distribution of COs along chromosomes at the megabase scale. **(A)** Total CO number per F2 plant. Each dot represents the CO number in an F2 plant. The average COs per F2 is indicated for each genotype. (**B**) Schematic representation of NRZ, LRZ1, and LRZ2 on the physical chromosome. The centromeric repeat array is indicated in grey. **(C)** CO distribution on all 5 chromosomes in wild type (grey), *spf2 (pink), sgo2 (green), ctf18 (blue),* and *cmt3 (purple)*. The y-axis corresponds to the CO frequency (cM/Mb) using a 1Mb window size and 50kb step size. The x-axis corresponds to physical distance in Mb in the Col-CEN v1.2 assembly. The gray shading indicates the centromeric *CEN178* repeats array, and the vertical dotted lines indicate the borders of the NRZ, LRZ1, and LRZ2. Asterisks indicate 2Mb non-overlapping intervals with a significantly different recombination rate than wild-type (corrected Mann-Whitney tests).

**Figure 3:**
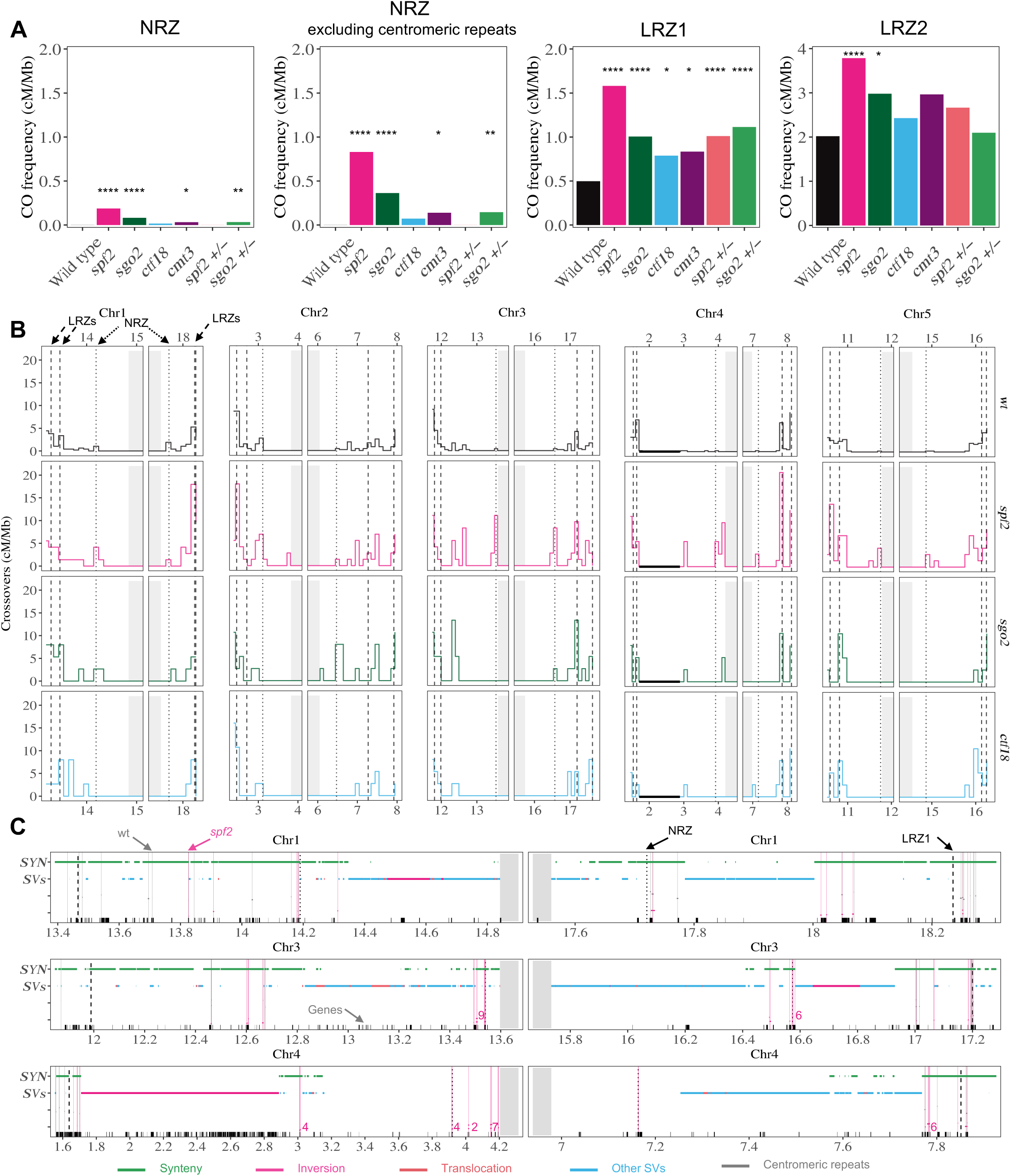
CO frequencies are increased in NRZ and LRZ in *spf2, sgo2, ctf18,* and cmt3. **(A)** CO frequency (cM/Mb) in NRZ, NRZ excluding the CEN180 repeat arrays, LRZ1, and LRZ2. The y-axis shows the CO frequency (cM/Mb), merging the five chromosomes. Stars correspond to Fisher exact tests (* p<0.05, ** p<0.001, **** p<10^-4^). Data on individual chromosomes is shown in Figure S4. (**B**) CO distribution in the centromeric region (NRZ and LRZs). CO frequencies are measured in non-overlapping 100kb windows in wild type (grey), *spf2 (pink), sgo2 (green), and ctf18 (blue)*. The y-axis represents CO frequencies (cM/Mb), and the x-axis physical position (Mb). The gray shading represents the centromeric repeat array, and the vertical dotted lines the NRZ, LRZ1, and LRZ2 borders. The black horizontal line on chromosome 4 indicates the well-described inversion between Col and L*er* genomes. (**C**) High resolution distribution of COs in NRZ and LRZ1 in wild type and *spf2*, on chromosomes 1, 3, and 4. Other chromosomes and genotypes are shown in Figures S6 and S7. The x-axis corresponds to physical distance in Mb. Note that the repeat array (gray shading) is truncated. Black ticks on the x-axis represent genes. Vertical lines indicate the position of COs detected in wild type (grey) and *spf2* (pink). When multiple COs occurred at overlapping positions, the counts are indicated. Horizontal bars indicate Col/L*er* syntenic regions (green), inversions (pink), translocations (red), and other structural variants (blue).

The global frequency of COs in *ctf18* and *sgo2* were similar to wild type (Mann–Whitney– Wilcoxon test, p>0.1), with ∼8 COs per F2 (Figure 2A). Our reanalysis of available datasets for *cmt3* (n=98), using the Col-CEN genome assembly, confirmed previous results with no significant difference from wild type (p=0.1) ^15^ (Figure 2A and S3). In contrast, we observed a significant increase in *spf2* with an average of 9.4 COs per F2 (p=3.10^-20^ compared to the pooled wild-type, p=0.003 compared to wild-type sister plants) (Figure 2A and S3).

Global CO distribution along chromosomes (Figure 2C) does not appear to be affected in *ctf18* and *sgo2* mutants, but for each of them, one pericentromeric interval had significantly more recombination than wild-type (2Mb intervals, corrected Mann-Whitney tests, p=0.022 and p=0.010, respectively) (Figure 2C). In *spf2,* COs are significantly increased in nine pericentromeric intervals, and in six intervals close to chromosome ends (Figure 2C), suggesting that SPF2 limits COs in the proximity of both centromeres and telomeres. In the *cmt3* mutant, the global CO distribution appears largely unaffected, except in two terminal intervals where frequencies are significantly lower, consistent with previous observations ^15^ (Figure 2C).

We then examined recombination patterns close to the centromere at higher resolution (100kb non-overlapping intervals, Figure 3). In the centromeric repeat arrays (gray-shaded), no COs were detected in either wild-type (4,730 gametes) or mutant lines (2,424 gametes), suggesting that meiotic COs do not occur within the core centromere (Figure 3A, 3B, S4 and S5B). In the NRZ, no COs are observed in the wild type, by definition (4,730 gametes, see above). In contrast, we observed COs within the NRZ in all four tested mutants, despite much smaller population sizes (22 COs in 362 F2 plants for *spf2,* 5/188 for *sgo2,* 1/188 *in ctf18, and* 1/96 *in cmt3*) (Figure 3A and S4). In the NRZ, *spf2* has a CO density of 0.2cM/Mb; If we exclude the CEN180 repeats arrays, the CO density reached 0.83 cM/Mb in the non-CEN180-NRZ (Figure 3A). Unlocking of COs is observed in NRZ of each chromosome in *spf2* (Figure 3B, Figure S4), and the effect is particularly marked in *spf2* in the NRZ of chromosome 4. Moving outwards, all the mutants revealed a significant increase in COs in LRZ1, the greatest effect being observed in *spf2* (Figure 3A, 3B and S4). From an average 1 cM per chromosome in wild-type LRZ1, it is increased by almost 2-fold in *sgo2*, *ctf18,* and *cmt3* and by more than 3-fold in *spf2* (Figure 3A). In *spf2,* the increase is significant for each chromosome and is particularly strong for chromosomes 3 and 4 (Figure 3B and S4). In LRZ2, CO frequency is also augmented in several mutants, the strongest effect being observed in *spf2* with a ∼2-fold increase (Figure 3 and S4).

We further investigated CO position at the fine scale at higher resolution, using deeper sequencing of *spf2*, *ctf18* and wild-type samples containing centromere-proximal COs ^75^. Fine mapping of the COs present in the NRZ and LRZ1 revealed that the COs cluster within genes in syntenic regions, and are absent within structural variants (e.g: inversions or insertions/deletions) (Figure 3C, S6 and S7).

In summary, *cmt3, ctf18, sgo2,* and s*pf2* mutations enhance CO formation in NRZ and LRZ1, where COs are strongly suppressed in the wild type. These peri-centromeric COs occur in syntenic regions and within the proximity of genes.

### Mutations in *SPF2* and *SGO2* have a co-dominant effect on centromere-proximal COs

Dominant mutation could facilitate the manipulation of CO rates in plant breeding schemes. We tested the effect of heterozygous mutation of *SPF2*, *CTF18*, *CMT3,* and *SGO2* on pericentromeric COs. In the CTL3.437.55 interval, we found a significant CO increase in *spf2/+* and *sgo2/+* but not in *cmt3/+* and *ctf18 /+* (Figure 4A, S5 and S9). We then analyzed genome-wide recombination in hybrids *spf2/+* and *sgo2*/+ using progeny sequencing, as described above. The average CO numbers per genome were similar to wild type (Figure 2A). In the NRZ, we did not detect COs in *spf2/+* (n=187 F2s) and found one in *sgo2/+* (n=188 F2s) (Figure3A and S4). In LRZ1, CO events were ∼2 folds more frequent than in the wild type in both *spf2*/+ and *sgo2*/+ (Figure 3A and S4). The *spf2* and *sgo2* mutations are thus co-dominant, the heterozygous mutant having more proximal CO than wild-types, but less than in homozygous mutants (Figure 4, Figure 3A).

**Figure 4:**
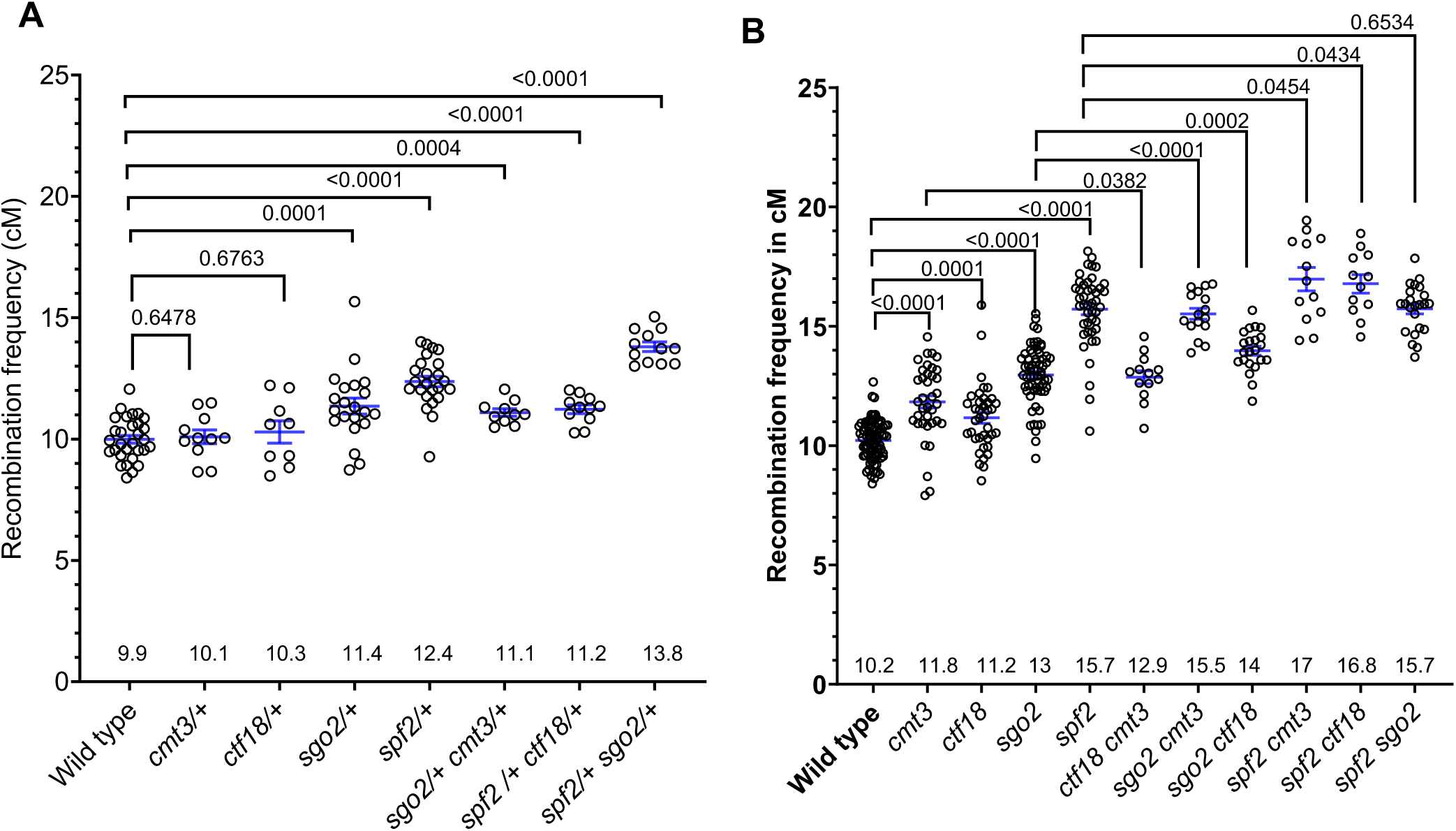
Cumulative effect of mutations on frequency of centromere-proximal CO. Recombination (cM) measurement in CTL.3.437.55, as in Figure 1A-B. **(A)** heterozygous mutants *ctf18/+, cmt3/+, sgo2/+, spf2/+*, and combinations. **(B)** Single and double homozygous mutants. These figures pool the data from several populations, and the data for Individual experiments are shown in Figures S9 and S10. Blue lines indicate the Mean of the recombination frequency +/- standard Error (SEM), and its value (cM) is given for each genotype. P-values are Mann-Whitney tests. Mutant alleles are *cmt3-11*, *ctf18-mp193, sgo2-1* and *spf2-3*.

### Parallel pathways repress COs in pericentromeres

Single mutations in *CTF18*, *SPF2*, *SGO2* and *CMT3* increase centromere-proximal COs, we aimed to test if they could have a cumulative effect, which would suggest that they belong to parallel pathways of CO repression. We measure CO frequencies in the CTL3.437.55 interval in the six possible double-mutants, together with single mutants and wild-type plants from the same segregating populations (Figure 4B and S10). We confirmed an increase of COs in each single mutant, compared to the wild-type, with *spf2* exhibiting the most pronounced effect (Figure 4B and S10).

All six double-mutants had increased COs compared to the wild type (Figure 4B and S10). To determine whether the effect on recombination was cumulative, we compared each double mutant to the highest single mutant in its respective combination. The double mutants with *cmt3 - ctf18 cmt3*, *sgo2 cmt3*, and *spf2 cmt3 -* showed significantly higher recombination than the corresponding single mutants (Figure 4B and S10). This suggests that CMT3 prevents centromere-proximal CO formation through parallel mechanisms relative to SPF2, CTF18 and SGO2.

Similarly, *spf2 ctf18* and *sgo2 ctf18* had significantly higher recombination than *spf2* and *sgo2*, suggesting that CTF18 suppresses COs independently from SPF2 and SGO2. However, *spf2 sgo2* recombination was not increased compared to *spf2*, the highest single mutant, implying that SPF2 and SGO2 may operate through a common pathway to prevent centromere-proximal COs (Figure 4B and S10). The two double mutants *spf2 cmt3* and *spf2 ctf18* had the highest recombination rate in CTL3.437.55 observed so far, an increase of ∼60% compared to the wild type.

### Enhanced pericentromeric CO does not appear to affect genome stability

Prior research suggested that centromere-proximal crossovers can cause chromosome missegregation at meiosis, notably in human^31,32^. We screened for aneuploid by analyzing the sequences of *ctf18*, *sgo2*, and *spf2* mutant progenies (F2 populations from Col/L*er* Hybrids), using unbalanced sequence coverage between chromosomes as the detection criteria ^75–77^. In wild-type (n=2365), *ctf18* (n=188), *cmt3* (n=96)*, sgo2* (n=188), *sgo2+/-* (n=188) and *spf2*+/- (n=187) we did not detect any aneuploids, suggesting that increasing crossovers at the centromere-proximal regions does not lead to higher aneuploidy rates. However, we found three trisomic plants in the progeny of *spf2* (0.8%, n=365), suggesting a slight genomic instability. Interestingly, the three trisomies were not associated with the presence of CO in the NRZ/LRZ1 (Figure S8), suggesting the chromosome missegregation events are not the consequence of proximal CO but an independent effect of the *spf2* mutation, presumably on cohesion and/or monopolar orientation (Singh et al, BioRxiv 2025).

Next, we assessed the fertility of mutants with elevated pericentromeric CO, as measured by the average number of seeds per fruit (Figure 5). None of the *cmt3*, *ctf18*, *sgo2*, and *spf2* single mutants showed a reduction in fertility compared to their respective sister wild-type plants. Similarly, the double mutants *ctf18 cmt3*, *sgo2 cmt3*, *spf2 cmt3,* and *sgo2 ctf18*, in which proximal CO are further elevated, the fertility was not significantly reduced compared to the wild type. However, fertility was reduced by 43% in *spf2 ctf18* and by 22% in *spf2 sgo2* compared to their sister wild-type plants (Figure 5). In *spf2 ctf18*, we observed a mild meiotic defect, with infrequent precocious separation of sister chromatids and segregation defect at meiosis II (Figure S11-12). In *spf2 sgo2,* we could not detect a clear meiotic defect (Figure S13-14). Interestingly, the observed reduction in fertility does not well correlate with increased proximal recombination: Combining *spf2* and *sgo2* mutation does not increase COs compared to the single mutants, but does reduce fertility (Figure 4B and 5); The highest levels of proximal COs were observed in *spf2 cmt3* and *spf2 ctf18* (Figure 4B), but fertility is affected only in *spf2 ctf18* (Figure 5). This suggests that the fertility reduction observed in some mutants is not the consequence of increased centromere-proximal COs but is caused by upstream or independent effects of the mutations.

**Figure 5:**
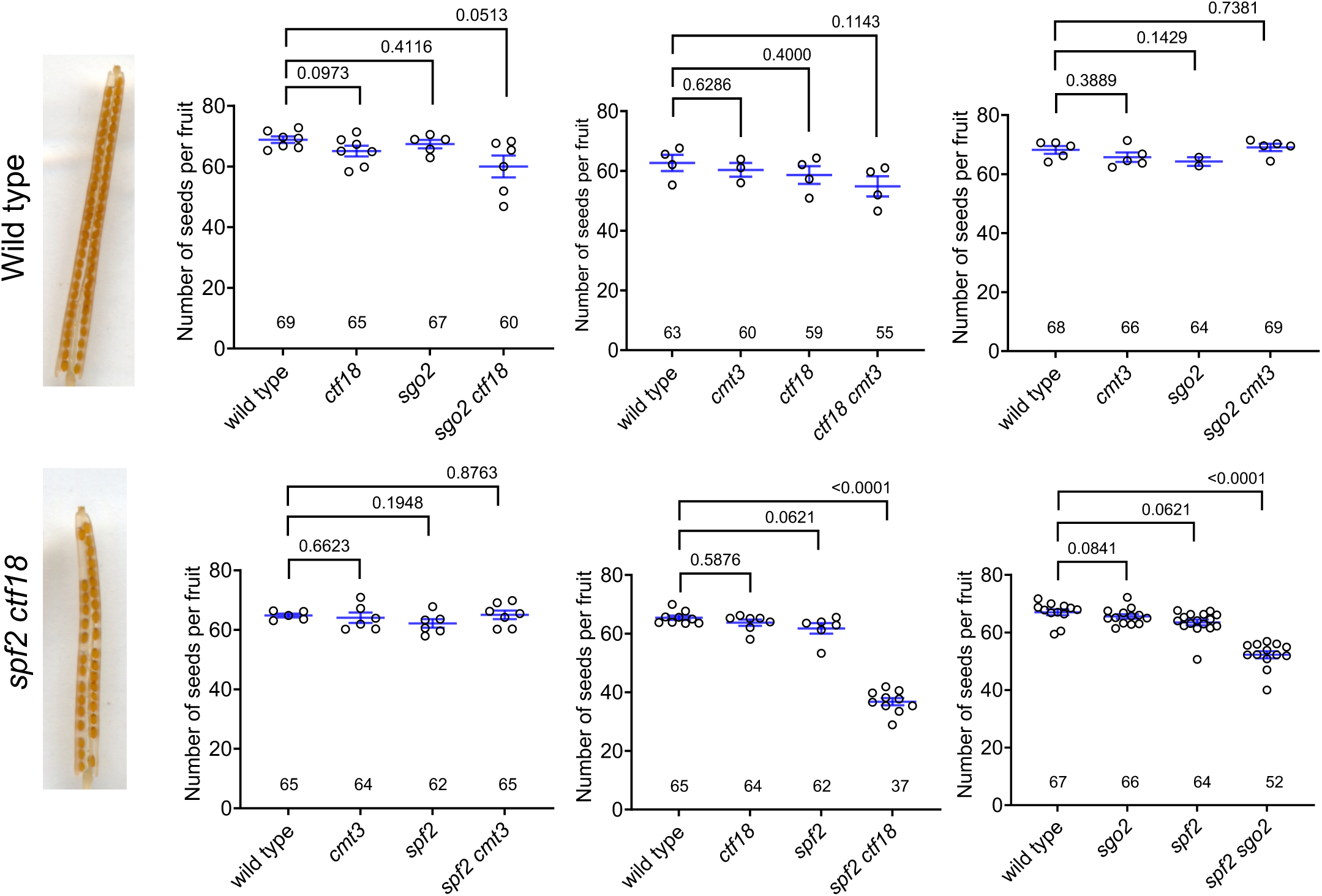
Fertility in mutants with elevated centromere-proximal Cos. Fertility was assessed by counting the number of seeds per fruit in 10 fruits on a minimum of three plants per genotype. Each dot represents the average number of seeds per fruit for an individual plant. Blue lines indicate the mean +/- Standard Error of the Mean (SEM), and its value is given. Representative pictures of wild-type and *spf2 ctf18* fruits are shown. Statistical comparisons are Mann-Whitney tests. Mutant alleles are *cmt3-11*, *ctf18-mp193, sgo2-1* and *spf2-3*.

## Discussion

The phenomenon of CO suppression near centromeres has been known for nearly a century, though the underlying mechanisms have remained elusive^18^. Suppression of centromere-proximal COs is an important limitation for plant breeding. In Arabidopsis, complete suppression of COs spans the 12 Mb centromeric repeat array and an additional 4 Mb of flanking chromatin. Furthermore, an additional 10 Mb flanking region (LRZ1) exhibits a strong reduction in COs.

This study shows that *CTF18* and *SGO2* play a role in restricting crossovers near centromeres in *Arabidopsis*, both being regulators of cohesin in eukaryotes. *CTF18* is a conserved replication factors that participate in sister chromatid cohesion establishment ^78^. Its role is conserved in Arabidopsis, but is not essential, making the growth and development of the *ctf18* single mutant indistinguishable from wild-type ^66^. Shugoshins are conserved proteins that protect centromeric cohesion during meiosis I ^69,79^. In Arabidopsis, both *SGO1* and *SGO2* contribute to cohesin protection at meiosis I, with SGO1 playing a prominent role, precocious loss of cohesion being observed in the *sgo1* but not in the *sgo2* single mutant ^70,71^. SGO1 and SGO2 also promote monopolar orientation of the kinetochore, with SGO2 playing a prominent role (Singh et al., BioRxiv 2025). Our result thus suggests that proximal COs are prevented by cohesins, and lessening cohesin loading or protection is sufficient to unlock proximal COs. Mutations in either SGO2 or CTF18 lead to an increase in centromere-proximal crossovers and combining the two mutations results in an even greater increase (Figure 4), possibly by reducing cohesin density further. In yeasts, cohesin regulation also contributes to CO suppression in centromere-proximal regions ^2,52^. The enrichment of cohesins at the centromeres could lead to CO suppression by regulating several steps of the meiotic recombination pathway: (1) A high cohesin density may prevent DNA double-strand break formation by limiting access to SPO11 and associated factors ^2^. (2) Preference for Inter-Sister Repair Over Inter-Homolog Repair – Cohesin enrichment near centromeres may promote sister chromatid-mediated DSB repair, thereby repressing COs. Studies in budding yeast support this model, showing that cohesins influence template choice between sister or homologous chromatids for DSB repair during vegetative growth ^53^. (3) Cohesins may favor the repair of recombination intermediates into non-COs rather than COs, as suggested by a higher non-COs/CO ratio in vicinity to centromere in humans ^26^.

We show that disruption of SPF2 leads to a large increase in CO formation in both the NRZ and the LRZs. SPF2 is a homolog of the yeast deSUMOylase Ulp2. Ulp2 regulates cohesins and condensins turnover by preventing the assembly of SUMO chains on SMC complexes ^72^. Thus, one possible function of SPF2 is the regulation of cohesins near centromeres, similar to *CTF18* and *SGO2*. This hypothesis is supported by several observations: (i) in combination with other mutations, *spf2* leads to premature loss of sister chromatid cohesion at meiosis (Singh et al., BioRxiv 2025); (ii) The amount of REC8 cohesin is reduced on metaphase meiotic chromosome in *spf2* (Singh et al., BioRxiv 2025); and (iii) combining *spf2* with *sgo2* did not lead to further CO increase in the proximal interval tested, suggesting that they act in the same pathway of CO suppression. SPF2 may directly regulate the activity of SGO2 in cohesion protection, a hypothesis supported by the known regulation of SGO turnover by SUMO in yeast ^80^. However, the observation that the mutation of *SPF2* leads to a higher frequency of proximal CO than *sgo2*, and the reduced fertility in *spf2 sgo2* suggests that SPF2 has additional targets. Possible additional targets could be SGO1 or cohesin subunits. The *spf2 ctf18* double mutant has more proximal COs than the single mutants, suggesting that they act via parallel pathways. One possibility is that they cumulatively regulate cohesins, which is suggested by the reduced fertility and meiotic defect of the double mutant. Alternatively, SPF2 may contribute to CO suppression independently of cohesins, possibly targeting kinetochores or centromeric chromatin ^52,81,82^. Lastly, SPF2 may also directly regulate the SUMOylation of the pro-CO recombination machinery, such HEI10 and MSH4, which were shown to be SUMOylated in yeast ^83,84^. Further investigation is required to test these hypotheses regarding the mechanism by which SPF2 limits proximal COs.

The DNA methylase CMT3 contribute to proximal CO suppression ^15^. Our data showed that *cmt3* has cumulative effects on recombination near centromeres (CTL3.347.5) with *sgo2, ctf18,* and *spf2,* indicating that DNA methylation and cohesin regulation independently contribute to centromeric CO suppression. One attractive scenario is that DNA methylation promotes closed chromatin and that cohesin acts at a different scale promoting folding of chromatin fibers and/or interacting with the recombination machinery.

In *Drosophila*, humans and yeasts centromere-proximal crossovers were associated with chromosome segregation defects ^2,31,32,34^. Despite the observed increase in centromere-proximal COs, fertility remains unaffected in *spf2, ctf18*, and *sgo2* mutants, indicating that enhancing centromere-proximal COs is possible without impacting chromosome segregation. We did not detect any aneuploids, in the progeny of *ctf18* and *sgo2* mutants and found three trisomic plants in the progeny of *spf2* (0.8%). However, the three trisomy were not associated with the presence of proximal CO, suggesting that the chromosome missegregation events are not the consequence of proximal COs but an independent effect of the *spf2* mutation, presumably related to cohesins. Among the mutant combination we tested, *spf2 ctf18* and *spf2 cmt3* showed the highest frequency of centromere-proximal COs. While *spf2 ctf18* has reduced fertility, *spf2 cmt3* does not. This suggests the reduced fertility of *spf2 ctf18* is not caused by centromere-proximal COs, but rather an independent effect likely related to weakened cohesin.

The combined challenges of food security and sustainable agriculture call for efficient plant breeding, which relies on meiotic recombination. Mutating generic anti-CO factors was shown to increase COs in model and crop species, but the effect is limited to chromosome arms and does not unlock pericentromeric regions that remain inaccessible to breeding ^85–91^. In this study, SPF2, CTF18, and SGO2 emerged as appealing targets to increase centromere-proximal COs in plants. All these proteins are conserved in crop plants and more distant eukaryotes. In Arabidopsis, *spf2* mutation showed the strongest increase in frequency of centromere-proximal CO, followed by *SGO2* and *CTF18*, making it the most promising candidate. It would be interesting to test all these factors in crops, for example tomato and cereals in which very large pericentromeric regions are devoid of crossovers ^4,5,23^, and contains gene of interests such as plant defense response genes^39^. The association observed in barley between natural genetic variation in the REC8 cohesin gene and crossover patterning suggests that the role of cohesin regulators in suppressing proximal COs is conserved in cereal crops^92^. Another line of research would be to combine generic anti-CO factors like *RECQ4* ^85,88^ with those enhancing centromere-proximal COs, testing for synergetic effects. The mutation of SGO2 and SPF2 showed to have a co-dominant effect in unlocking pericentromeric COs, which may facilitate their integration in plant breeding programs ^93^. Unlocking CO in pericentromeres would allow mapping of QTLs and separation of beneficial traits from undesirable alleles in so-far inaccessible region ^42,44^.

In conclusion, SPF2, SGO2, and CTF18 contributes to the prevention of centromere-proximal COs. Discovering these genes in Arabidopsis advances our understanding of centromere-proximal recombination suppression. The capacity to increase centromere-proximal COs without affecting fertility opens new perspective for crop improvement.

## Materials and Methods

### Plant materials and growth conditions

*Arabidopsis* plants were grown in a greenhouse or growth chamber under controlled long-day conditions (16 h day/8 h night, 20 °C). The Arabidopsis lines used in this study were as follows: Fluorescent lines to measure recombination: CG437 (CS71169) and CR55 (CS71173) ^56^. Mutant lines used in the study: *cenpc* (*cenpc-mp221*) Col-0 (Singh et al., BioRxiv 2025); *ctf18* (*ctf18-mp193*) Col-0 (Singh et al., BioRxiv 2025); *smc1* (*smc1-mp172*) Col-0 (Singh et al.); *smc3* (*smc3-mp226*) Col-0 (Singh et al., BioRxiv 2025); *zyp1 (zyp1-1)* Col-0 and *zyp1 (zyp1-6*) L*er* ^65^; *cmt3* (*cmt3-11*-SALK_148381) Col-0 ^15^; *smc1* (FLAG024C06, line DPF3) Wassilewskija (WS))^94^; *sgo2-1* (line 34303) ^70^; *spf2-3* (SALK_140824) ^95^. Plants were genotyped by PCR with an annealing temperature of 58°C (primers in Table S1).

### Recombination measurements using Fluorescent Tagged Lines (FTLs)

To measure recombination, we utilized fluorescent tagged lines (FTLs) developed in the Col-0 accession ^56^. We crossed a plant carrying the T-DNA line CG437 (CS71169) with a GFP (green fluorescent protein) marker to another plant carrying the T-DNA line CR55 (CS71173) with a dsRed fluorescent protein marker. Both fluorescent proteins are expressed in seeds under the control of the *NapA* promoter. These T-DNA markers flank the centromere on chromosome 3. We obtained F1 plants that displayed both red and green fluorescence in trans. Our goal was to have these colors in cis configuration to measure recombination between the two markers. To achieve this, we grew the F2 progeny and identified plants that were homozygous for both the red and green T-DNA markers, indicating that the markers were in cis configuration. We also selected plants that were homozygous for one marker and heterozygous for the other, as they also had the markers in *cis* on one chromatid. These selected plants were used in further crosses with candidate genes to measure recombination.

We then crossed the candidate gene with the cis configuration line, named CTL3.347.55, to obtain F1 seeds. These seeds were grown into plants heterozygous for both the candidate gene and the red and green T-DNA markers. These plants were self-pollinated to obtain F2 seeds. From the F2 seeds, we preselected those that were heterozygous for both T-DNA markers, as heterozygosity is required to visualize recombination events. We then genotyped the F2 plants to identify those that were homozygous mutants for the candidate gene, as well as their wild-type siblings. We allowed these plants to self-pollinate to produce F3 seeds, in which recombination was measured for both the homozygous mutant and sister wild-type plants.

For recombination analysis, we employed two different methods: one using CellProfiler, a previously established method ^96^, and another established in the group using Fiji image analysis software, coupled with a pipeline for data analysis using Perl and R described in the following publication ^97^. Seed preparation differed between the two methods. For CellProfiler, seeds were spread evenly on a black surface using a microscopic slide to prevent overlap. For the Fiji method, seeds were placed on 1% agar in small Petri dishes to ensure separation and prevent overlap. For figure 1B and S2 C-G, recombination frequency was measured using the Fiji image analysis software method and by placing seeds on 1% agar. For Figure 4 and S9, S10, recombination frequency was measured using the CellProfiler method and by using dry seeds.

In both methods, three images were acquired for each sample: (i) brightfield, (ii) UV through a dsRed filter, and (iii) UV through a GFP filter, using a fluorescent stereomicroscope. CellProfiler/ Fiji image analysis (version 2.1.1) was used to identify seed boundaries in the micrograph images and assign fluorescence intensity values for RFP and GFP to each seed. Genetic distance was calculated using the formula:

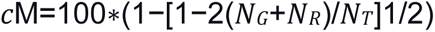

where N_G_ is the number of green-alone fluorescent seeds, N_R_ is the number of red-alone fluorescent seeds, and N_T_ is the total number of seeds counted ^96^. The Mann-Whitney test was used for statistical comparison between samples.

### CRISPR-Cas9 mutagenesis of *AtSGO2, AtCTF18* and *AtSPF2*

All guide RNAs were designed based on the Arabidopsis Columbia-0 genome and verified for specificity against the Landsberg genome using the CRISPR-P 2.0 tool http://crispr.hzau.edu.cn/CRISPR2/ with the U6 promoter. Two guides were selected at both the 3’ and 5’ ends, excluding UTRs (Table S1, Figure S1). The guides, rich in GC content, were 19 bp long, lacked repeats, and had the PAM sequence in the reverse complementary orientation. The guides were cloned into the plCH86966 plasmid using CloneAmp at 65°C. In the first reaction each guide (in total four guides per gene) was assembled into a vector that finally comprises of AtU6 promoter and guideRNA expression cassette. The vector was transformed into E. coli, purified, and Sanger-sequenced to confirm key features (restriction sites, the anchor site of each plasmid (reverse complementary) and the guide). Cloning of the guide RNAs into the final Cas9-containing binary vector was carried out using the MoClo Golden Gate cloning toolkit ^98^. In the second reaction, all four plasmids containing guide RNA were subcloned into the final binary vector, assembled with the intron-containing Cas9 ^99^ expression cassette, Fast-Red expression cassette (that allowed selection of Cas9 transformants by selecting for fluorescent red seeds) and linker, transformed into competent cells, selected with kanamycin, and verified by restriction digestion. The complete plasmid structure was confirmed by Nanopore sequencing https://www.plasmidsaurus.com/, and the plasmids were transformed into Agrobacterium (GV3101), selected with appropriate antibiotics, and verified by PCR. Transformed bacteria were selected from an overnight preculture in LB media containing gentamycin, kanamycin, and rifampicin. These were then incubated in fresh culture for approximately 5 hours until reaching an optical density of 0.5-0.6. Bacteria were centrifuged (4000 g for 10 minutes), resuspended in a freshly prepared 5% sucrose solution with 20 μl of Silwet L-77, and used to dip young flower buds of about 16 Columbia and Landsberg plants per batch for 15 seconds. The plants were then covered with dark plastic bags overnight. Transformed seeds (T1) were selected based on red fluorescence, Cas9 amplification by PCR was performed on T1 plants, mutations confirmed by PCR and by Sanger sequencing. T1 plants were selfed and in the T2 generation, we selected for mutants that are Cas9 free both based on selecting non-fluorescent seeds and validating the T2 plants by PCR, genotyping for the mutation in the gene and validating via sequencing. If the T2 plant is homozygous for mutation then the plant was backcrossed to Ler before crossing it to *sgo2-1, ctf18-mp193, spf2-2* alleles in Col-0. If the T2 Ler plant was with heterozygous mutation then the plant was directly crossed to its allele in Col-0. This way we produced, *ctf18-mp193/ ctf18-del* at G181, *spf2- 3/ spf2-del* at L90 and *sgo2-1/ sgo2-del* at K74 Col/Ler F1 hybrids. These F1 were validated by PCR for the mutations both in Col and Ler background and were selfed to produce F2 progeny which were sequenced (see below), to obtain genome-wide CO landscape.

### Genome-Wide sequencing

From the set of 334,680 high-quality Col/L*er* SNPs created from previous study ^8^, we further filtered those SNPs that distributed outside the 0.05-quantile and 0.95-quantile of total read number, Col allelic ratio (Col read number divided by total read number) and L*er* allelic ratio (Ler read number divided by total read number), homozygous Col allelic ratio (number of samples with 0/0 genotype divided by the total number of genotyped samples), heterozygous Col allelic ratio (number of samples with 0/1 genotype divided by total number of genotyped samples), homozygous Ler allelic ratio (number of samples with 1/1 genotype divided by total number of genotyped samples) and the total number of genotyped samples. Finally, a set of 234,744 of golden SNPs were kept for subsequent analysis.

In this study, we generated and sequenced F2 plants derived from F1 plants segregating for *spf2* (number of sequenced F2 plants: wild type n=284, *spf2*+/- n=187 and *spf2*-/- n=365), or *ctf18* (wild type n=188 and *ctf18*-/- n=188), or *sgo2* (*sgo2*+/- n=188 and *sgo2*-/- n=188). The sequencing dataset of the population of *cmt3* (wild type n=350 and *cmt3*-/- n=96) was downloaded from previous study ^15^. We also included F2 wild-type samples from Rowan et al; 2018. These data were analyzed/reanalyzed with the genome assembly Col-CEN v1.2. ^3^. The golden SNPs were genotyped by using inGAP-family ^100^, and then a sliding window-based method was used for predicting COs, using a window size of 50 kb and a step size of 25 kb ^75,76,100^. We further filter false close double COs that commonly present across samples, which are caused by mis-genotyping.

For high resolution mapping, we selected samples with centromere-proximal COs and re-sequenced with high depths, including samples from the *spf2* population (wild type n=23, *spf2*+/- n=18, *spf2*-/- n=101) and the ctf*18* population (wild type n=11, *ctf18*-/- n=20). We also included proximal COs from wild-type F2, male and female BC1 from previous study ^8,74,75^. We manually examined centromere-proximal COs with a high resolution.

The non-recombining zones (NRZs) were defined as the contiguous regions containing the main CEN178 arrays where CO are completely absent in wild type. The external borders of the NRZ are thus defined by the closest CO to the centromere detected in wild-type (19,167 COs from 4,730 gametes). Moving outward, we defined the LRZ1 (Low Recombining Zone), spanning 0.5 cM on each side flanking the boundaries of NRZs and the LRZ2 (Low Recombining Zone, defined by the position of the 24^th^ CO on each side), spanning 0.5 cM on each side flanking the LRZ1 (defined by the position of the 47^th^ CO on each side).

### Chromosome spreads preparations

Young inflorescences were harvested from Arabidopsis plants when they were 5-6 weeks old and inflorescences were fixed in Carnoy’s fixative (3:1 absolute ethanol: glacial acetic acid). Refresh the fixative until the solution is clear. The buds in fixative can be stored at 4°C. Flower buds roughly 0.5mm in size were isolated from inflorescences and washed in 10mM citrate buffer at pH4.5, then incubated in digestion mix (0.3% (w/v) Pectolyase Y-23 (MP Biomedicals) 0.3% (w/v) Driselase (Sigma) and 0.3% (w/v) Cellulase Onozuka R10 (Duchefa) for 2-3 hours at 37°C. Two to three digested buds were transferred to a clean slide and transformed into a fine cell suspension with a bent dissection needle. 10uL of 60% glacial acetic acid was added to the mixture and stirred gently at 45°C on a hotplate for 2 minutes. After one minute, one more 10ul drop of 60% glacial acetic acid was added to the slide and stirred gently. Stirring likely facilitates in removing the cytoplasm as much as possible. Cells were fixed on the slide by adding ice-cold 3:1 Carnoy’s fixative first around the droplet and then directly flushed on the drop. Excess of fixative removed by tilting the slide onto the beaker. Slides were left to dry tilted at room temperature, then 12uL of DAPI in anti-fade mounting media (Vectashield) was applied to the slide and a coverslip was added. Chromosomes were visualized using a Zeiss Axio Observer epifluorescence microscope.

## Acknowledgments

This work was supported by core funding from the Max Planck Society, and Alexander von Humboldt Fellowships to Joiselle Blanche Fernandes and Qichao Lian. Divyanshu Sahu is a recipient of the predoctoral fellowship HORIZON-MSCA-2021-COFUND-01 rePLANT-GA101081581 Co Funded by the European Union. Views and opinions expressed are however those of the author(s) only and do not necessarily reflect those of the European Union or the EUROPEAN RESEARCH EXECUTIVE AGENCY (REA). Neither the European Union nor the granting authority can be held responsible for them.

## Author contributions

RM lead the project. RSG, JBF, SD, DS, LCP and AK produced the genetic material, and conducted and analyzed biological experiments. Q.L. analyzed the sequencing data to assess recombination and aneuploidy. RSG, JBF, QL and RM wrote the manuscript, with inputs from other authors.

## Competing interests

The authors declare the following competing interests: A patent was filed by the Max Planck Society on the use of CTF18, DCC1, SGO2 and SPF2 to manipulate meiotic recombination in plants, with RM, RSG, JBF, SD, and QL listed as inventors (EP25184806.5. 24.06.2025).

---

**Figure S1:**
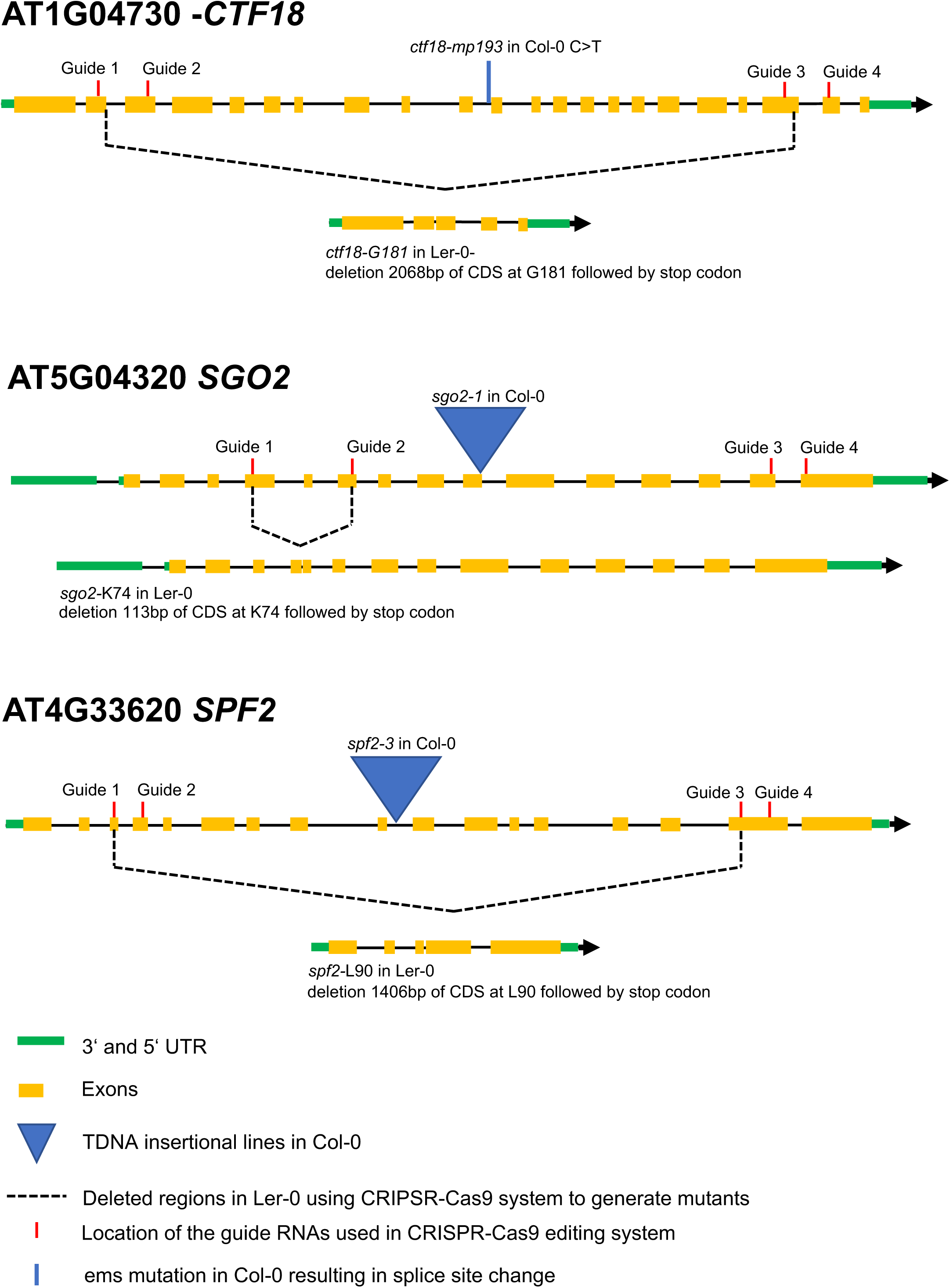
Gene and mutations used in this study.

**Figure S2.**
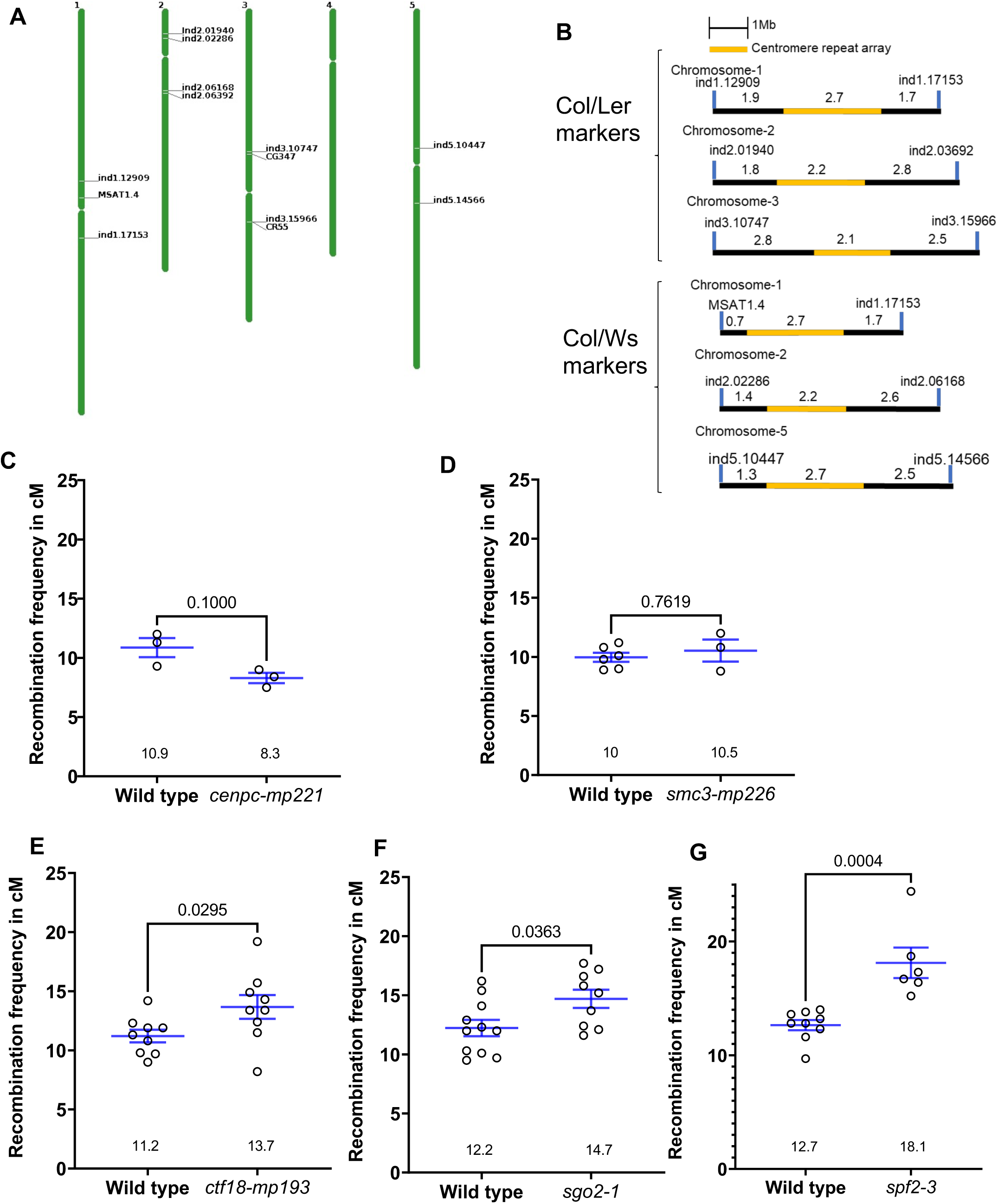
Measurement of recombination in individual populations. (A) Graphical representation of the genomic position of the markers used to measure recombination in CTL.3.437.55, Col/Ler and Col/Ws. (B) Schematic representation of the physical intervals defined by the markers used to measure recombination in Col/Ler and Col/Ws hybrids. Physical sizes are indicated in Mb. (C-G) Related to Figure 1B. Measure of recombination frequency in centimorgans (cM) in the CTL.3.437.55 interval. Each dot represents a measurement on ∼500 seeds. Blue lines indicate the Mean of the recombination frequency +/- standard Error (SEM), and the value (cM) is given for each genotype. Each mutant is compared to wild type from the same segregating population (sister plants). Statistical tests are Mann-Whitney tests.

**Figure S3.**
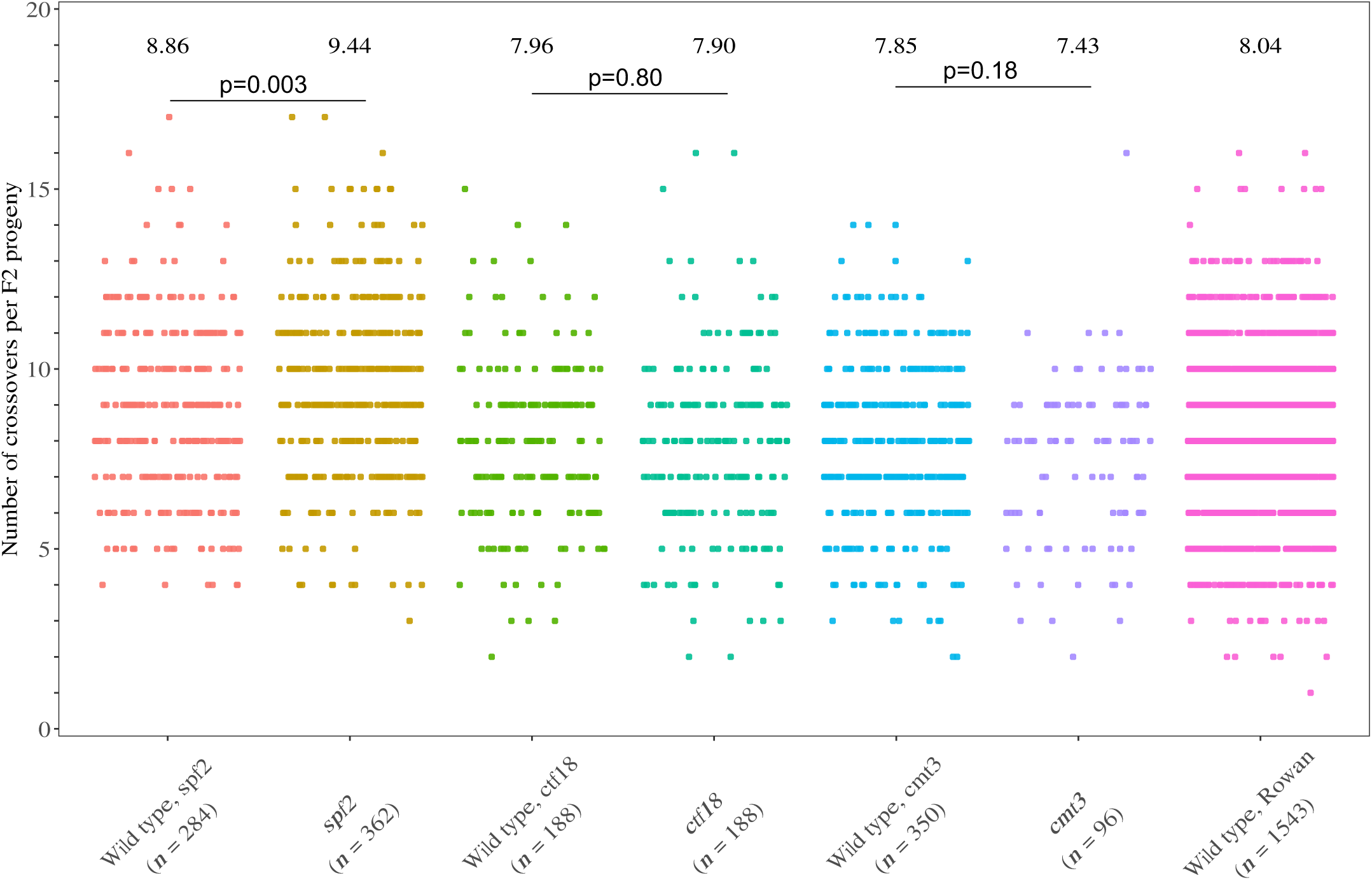
Total CO number per F2 in wild type and mutants. Related to Figure 2. Total CO number per F2 in wild type and mutants. Each dot represents the CO number in an F2 plant, and the average COs per F2 is indicated for each genotype. Mutants *spf2* and *ctf18* are compared to wild type from the same population (sister plants). The *cmt3* and corresponding wild-type data are from Underwood et al. ^15^. The large wild-type population is from Rowan et al ^101^. P values are Mann-Whitney tests.

**Figure S4:**
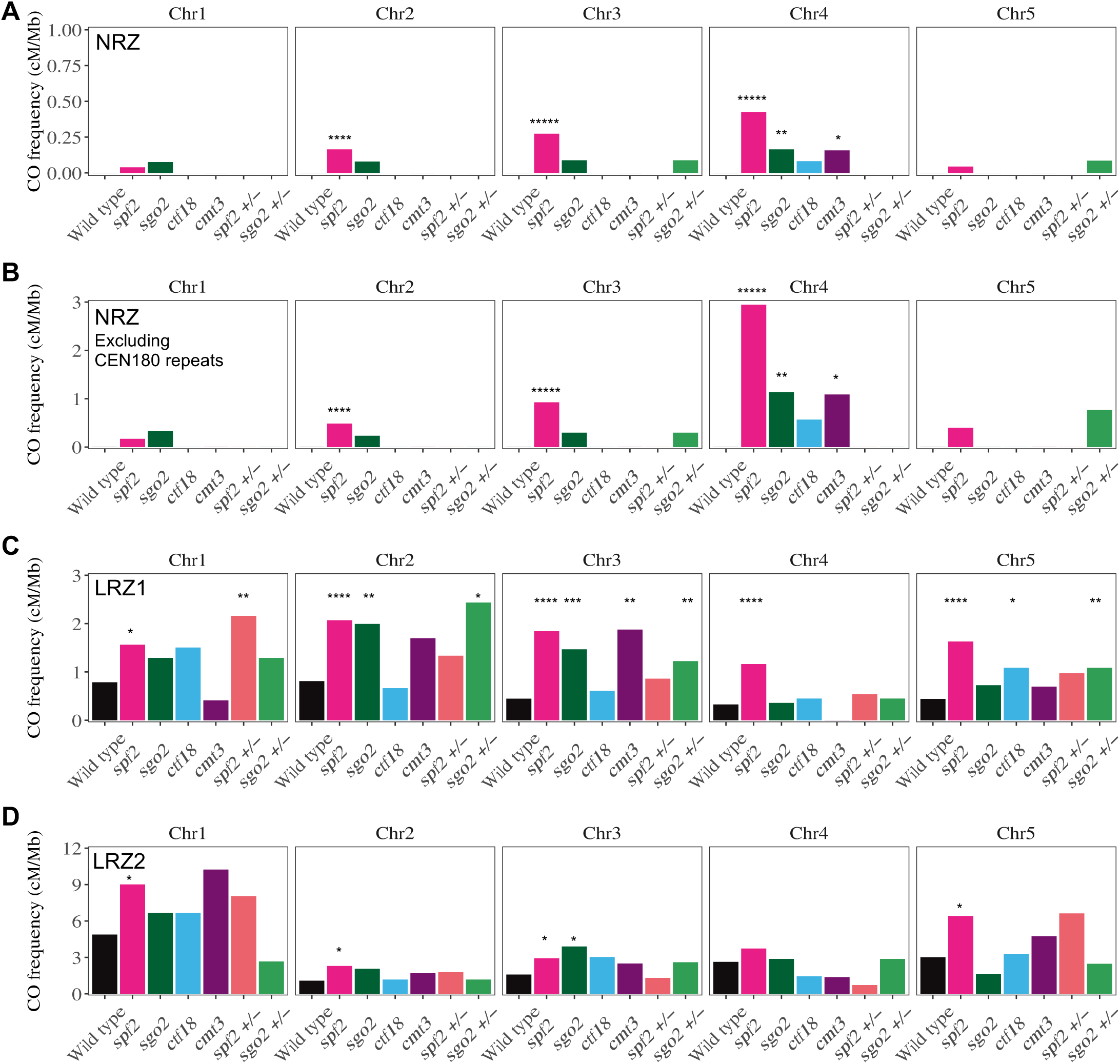
CO frequency in NRZ, LRZ1 and LRZ2 in wild type and mutants. Related to Figure 3A. Y-axis indicates the CO frequency in (A) NRZ, (B) NRZ excluding the CEN180 repeat array, (C) LRZ1 and (D) LRZ2, measured in cM/Mb for individual chromosomes. Stars correspond to Fisher exact tests on counts of samples with/without a CO in the considered zone compared to wild-type (* p<0.05, ** p<0.001, **** p<10^-4^)

**Figure S5.**
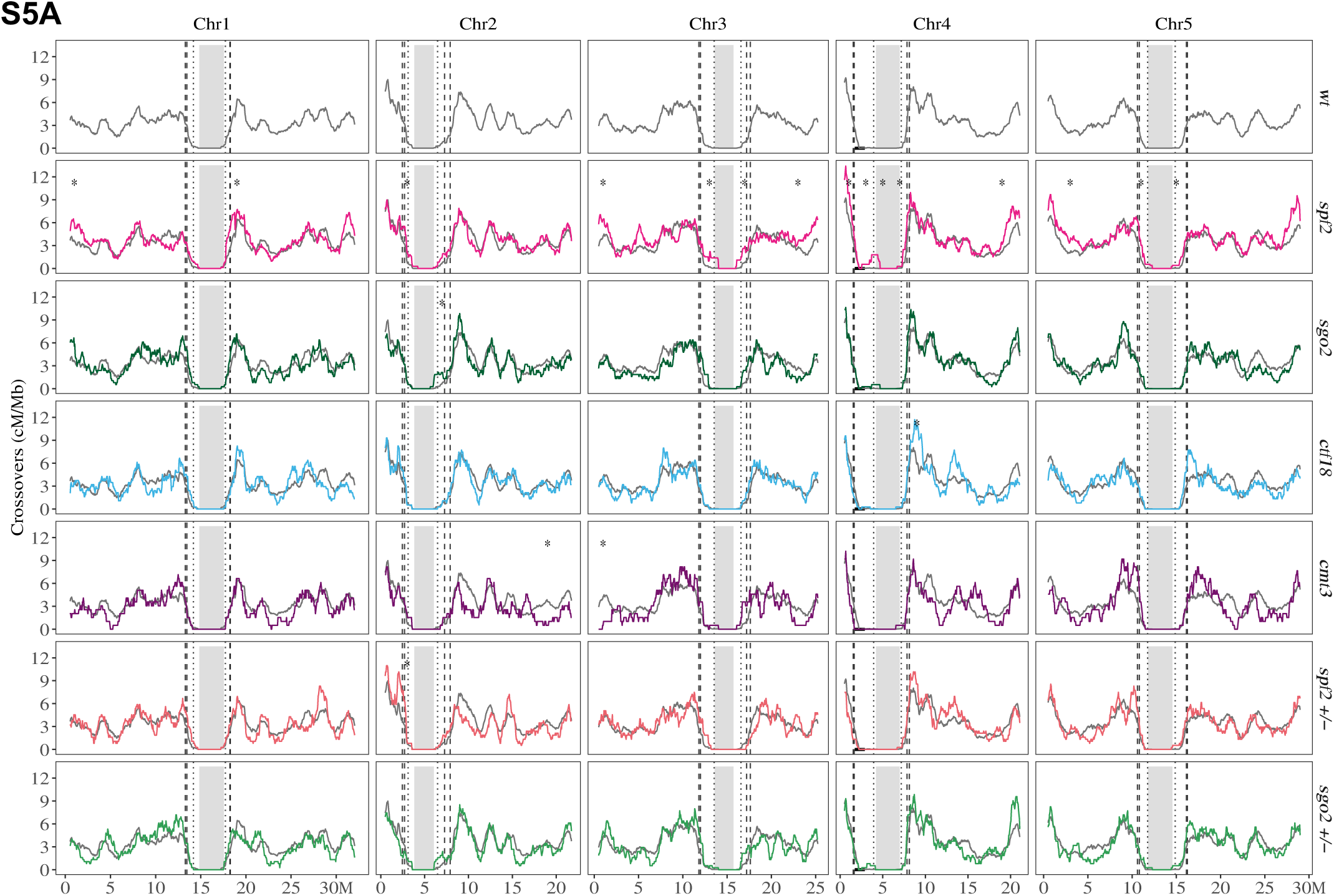
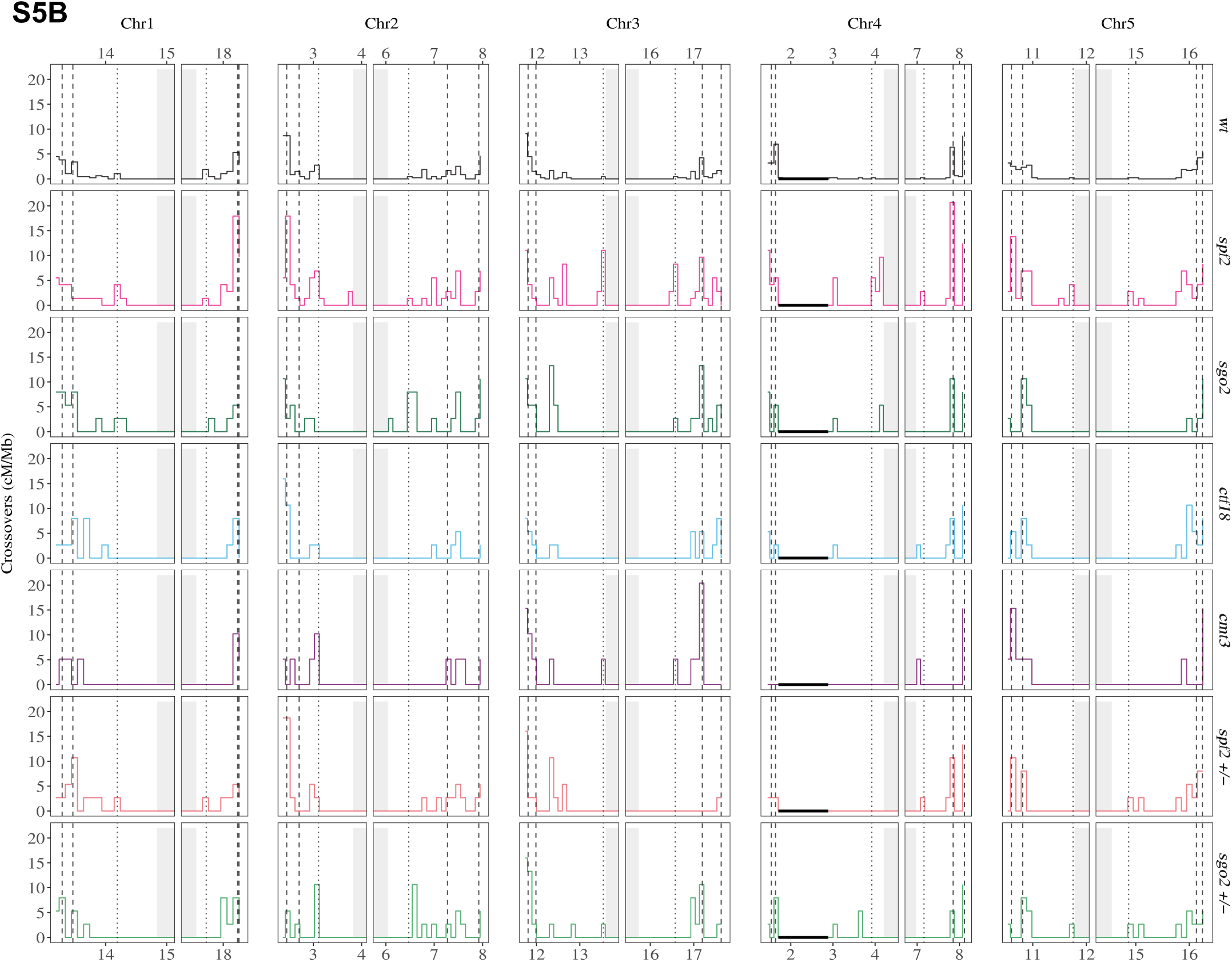
CO distribution in *spf2/+* and *sgo2*/+. (A) Genome-wide CO distribution on all 5 chromosomes in wild type (grey), *spf2 (pink), sgo2 (dark green), ctf18 (blue),* and *cmt3 (purple)* mutants, *spf2*+/- (orange) and *sgo2*+/- (light green). Y-axis corresponds to crossovers (cM/Mb) using a 1Mb window size and 50kb step size. The X-axis corresponds to physical distance in Mb (Megabases) in the Col-CEN v1.2 assembly. The gray shading indicates the centromeric *CEN178* repeats array and the vertical dotted lines indicate the borders of the NRZ, LRZ1 and LRZ2. Asterisks indicate 2Mb intervals with a significantly different recombination rate compared to the wild type (Mann-Whitney tests). (B) Crossover distribution in the centromeric region (NRZ and LRZ). Crossover frequencies are measured in non-overlapping 100kb windows. Y-axis represents crossover frequencies (cM/Mb), and X-axis corresponds to physical distance in Mb (Col-CEN). The gray shading represents the centromeric repeat array, and the vertical dotted lines the NRZ, LRZ1, and LRZ2 borders. The black horizontal line on chromosome 4 indicates the well-described inversion between Col and L*er* genomes.

**Figure S6.**
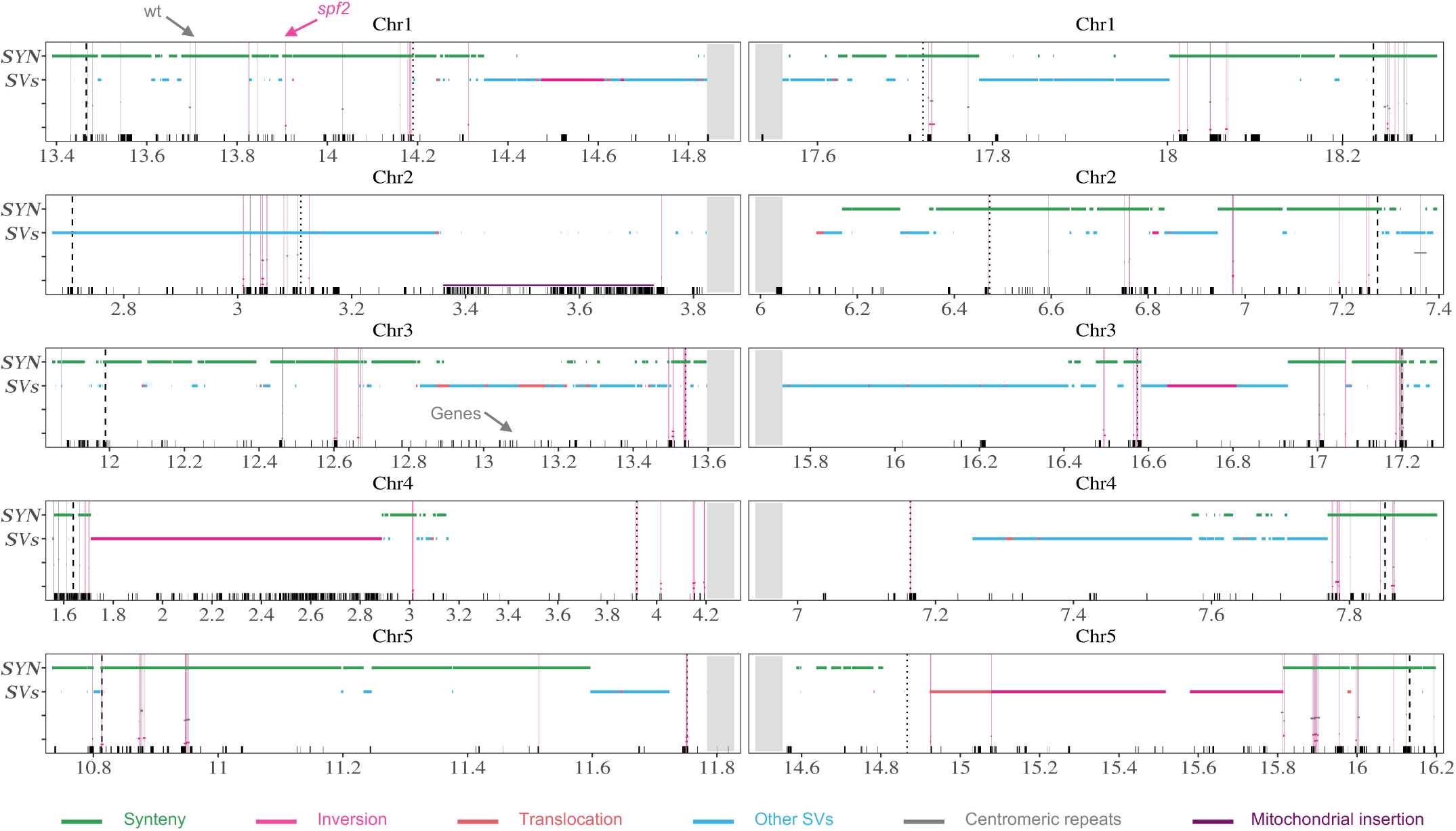
High resolution distribution of COs in NRZ and LRZ1 in wild-type and spf2. Related to Figure 3C. The X-axis corresponds to physical distance in Mb. Vertical lines indicate the position of COs detected in wild type (grey) and *spf2* (pink). When multiple COs occurred at overlapping positions, the counts are indicated. Grey shading shows the centromeric repeats (truncated in the middle). Horizontal bars indicate Col/L*er* syntenic regions (green), inversions (pink), translocations (red), a mitochondrial insertion (purple) and other structural variants (blue). Black ticks on the X-axis marks genes.

**Figure S7.**
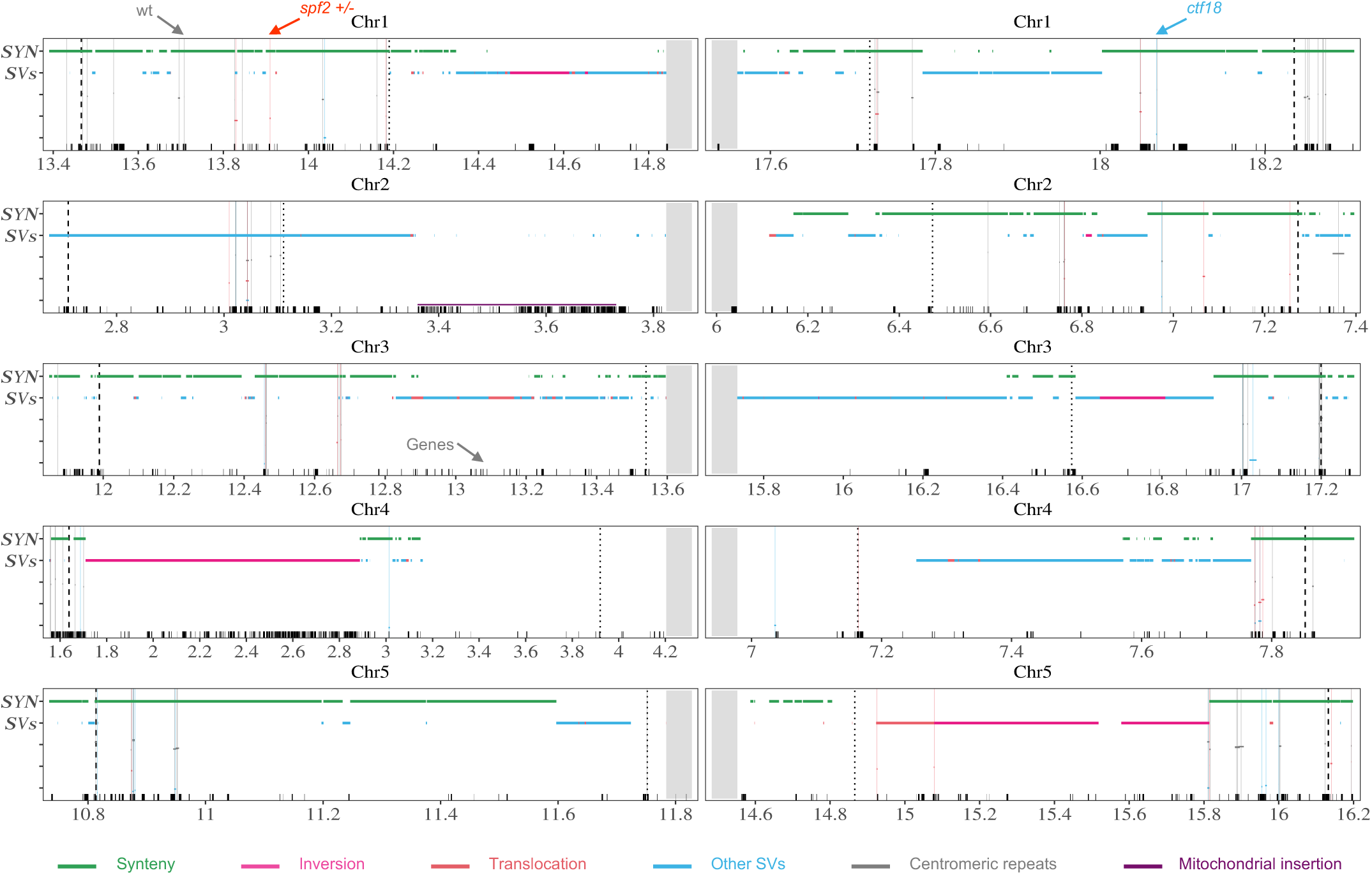
High resolution distribution of COs in NRZ and LRZ1 in *spf2+/-* and *ctf18*. Related to Figure 3C. The X-axis corresponds to physical distance in Mb. Vertical lines indicate COs detected in wild type (grey), *spf2+/-* (orange) and *ctf18* (Blue). When multiple COs occurred at overlapping positions, the counts are indicated. Grey shading shows the centromeric repeats (truncated in the middle). Horizontal bars indicate Col/L*er* syntenic regions (green), inversions (pink), translocations (red), a mitochondrial insertion (purple), and other structural variants (blue). Black ticks on the X-axis mark genes.

**Figure S8.**
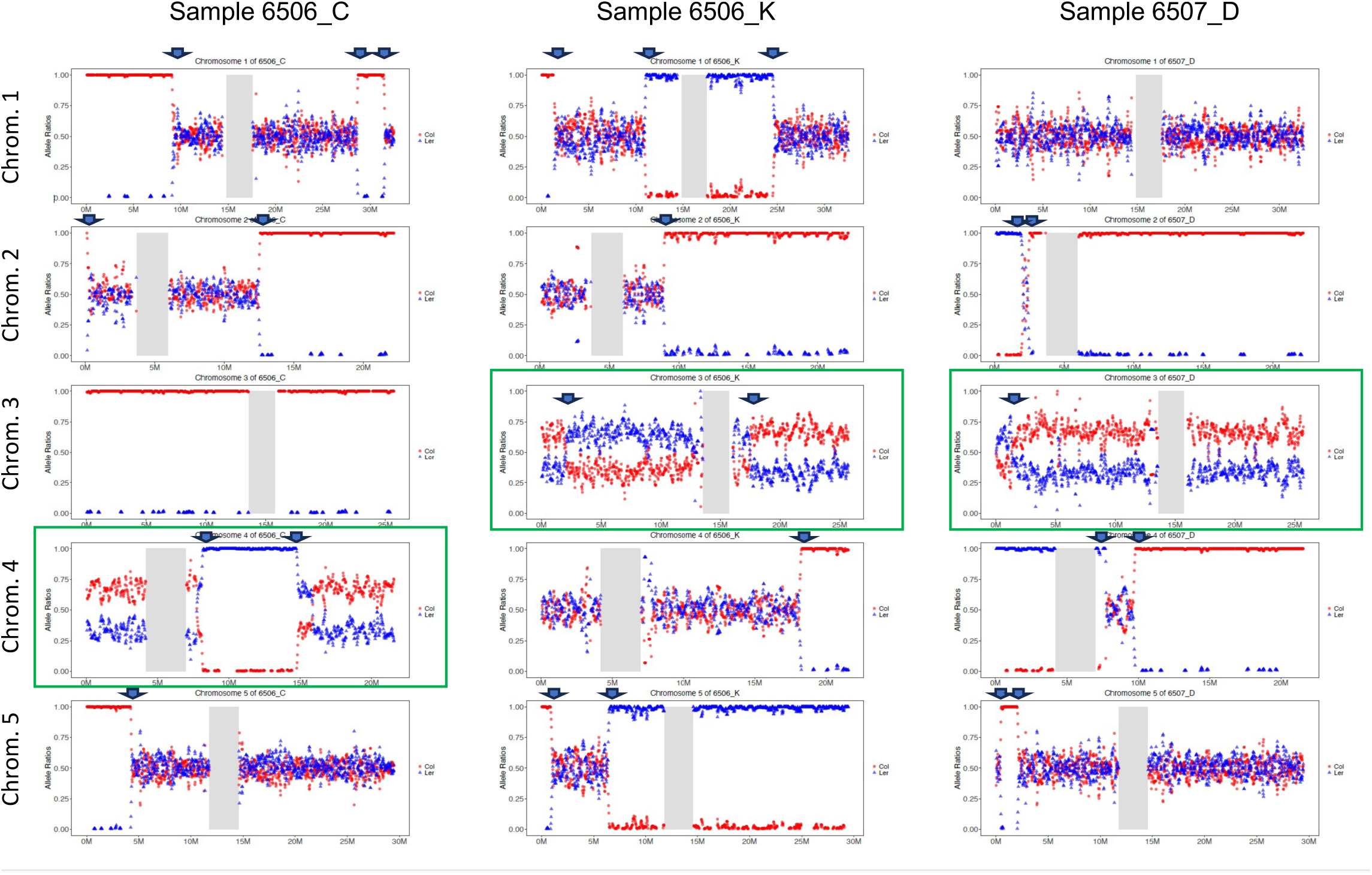
CO analysis in trisomic samples. The plots show the Col/L*er* allele frequency (Red/Blue) along the five chromosomes on the three aneuploid samples detected in the *spf2* progeny. Sample 6506_C is trisomic for chromosome 4, 6506_K and 6507_D are trisomic for chromosomes 3, as shown by higher sequence coverage and 2:1 allelic ratio. CO sites are indicated by arrows.

**Figure S9.**
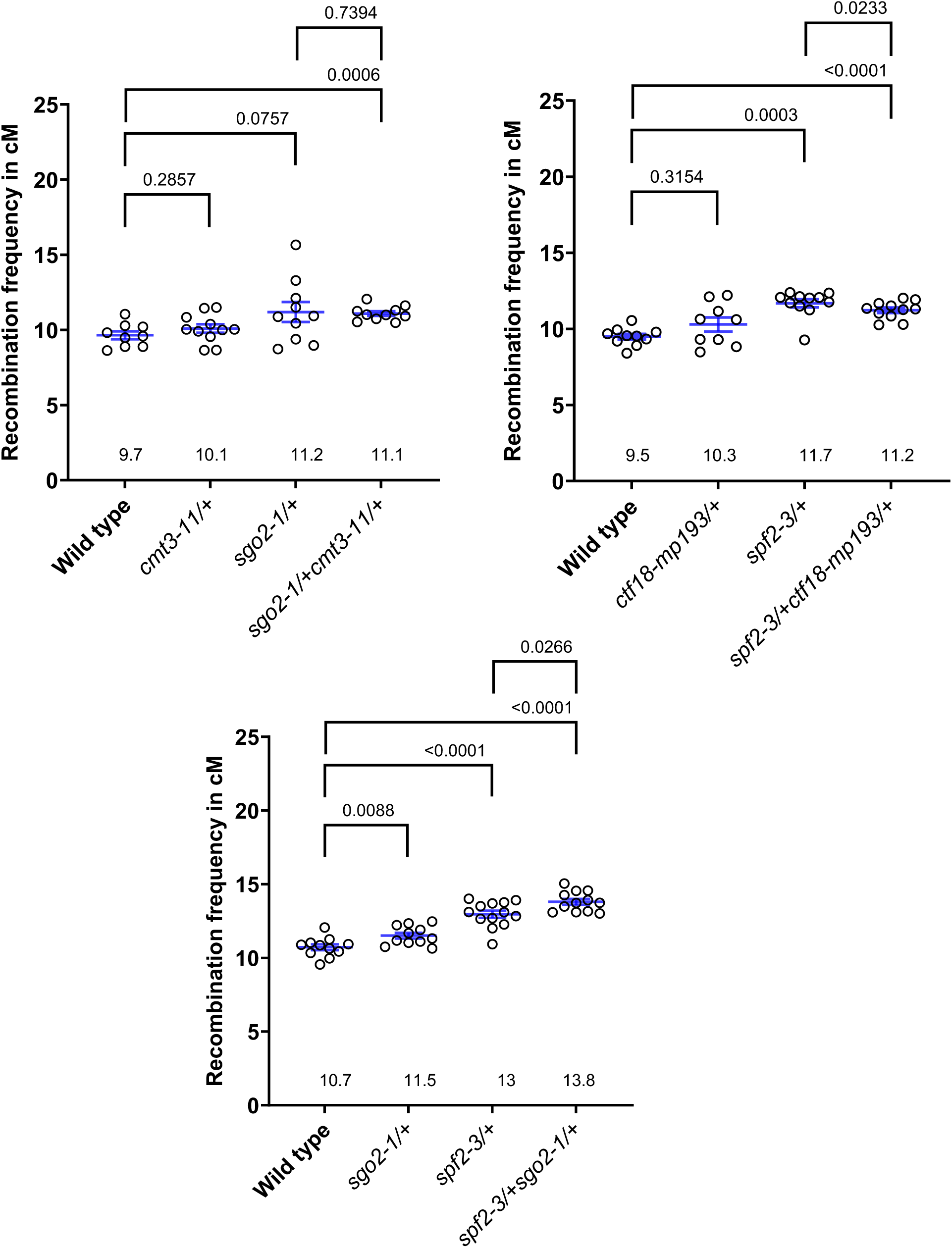
Measurement of recombination in CTL.3.437.55 in individual populations of heterozygous mutants. Related to Figure 4A. Measure of recombination frequency in centimorgans (cM) in the CTL.3.437.55 interval. Each dot represents a measurement on ∼500 seeds. Blue lines indicate the Mean of the recombination frequency +/- standard Error (SEM) and the value (cM) is given for each genotype. Single heterozygous, double heterozygous mutants and wild types from the same segregating population (sister plants) are compared. Statistical tests are Mann-Whitney tests.

**Figure S10:**
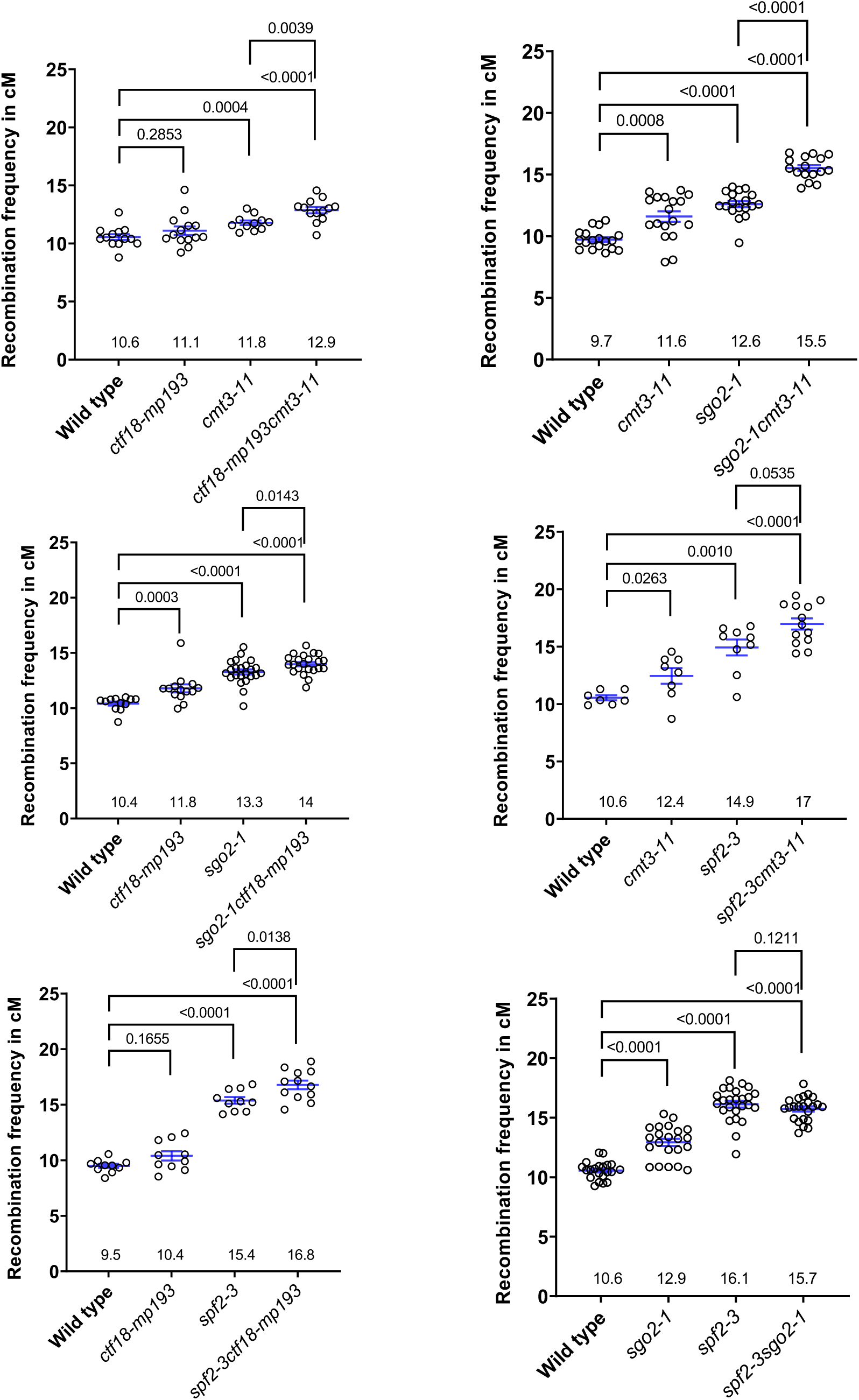
Measurement of recombination in CTL.3.437.55 in individual populations of double mutants. Related to Figure 4B. Measure of recombination frequency in centimorgans (cM) in the CTL.3.437.55 interval. Each dot represents a measurement on ∼500 seeds. Blue lines indicate the Mean of the recombination frequency +/- standard Error (SEM) and the value (cM) is given for each genotype. Each double-mutant mutant is compared to single mutants and wild types from the same segregating population (sister plants). Statistical tests are Mann-Whitney tests.

**Figure S11.**
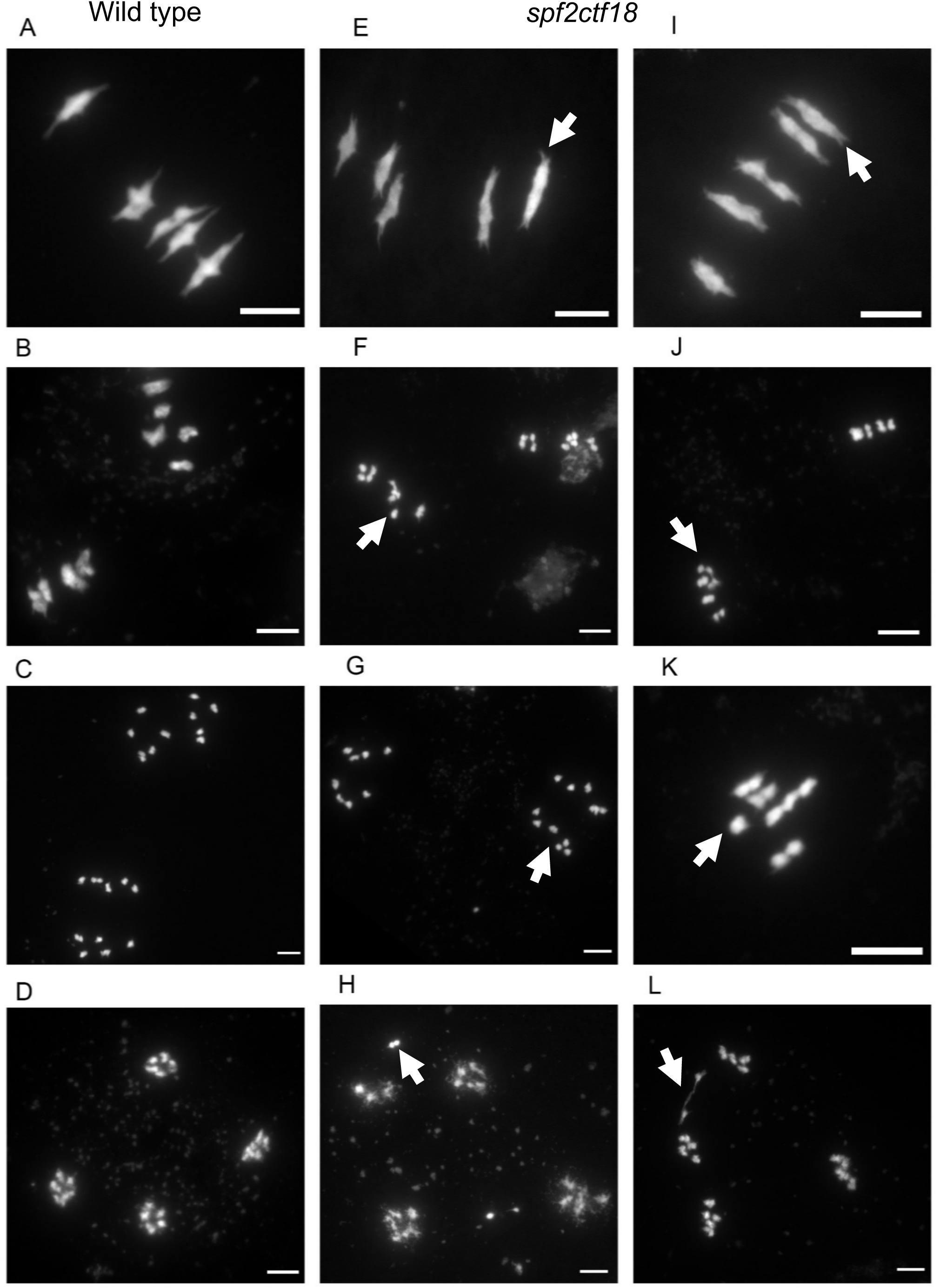
Meiotic Chromosome spreads in wild type and *spf2 ctf18*. Meiotic chromosome spreads were visualized using DAPI (4′,6-diamidino-2-phenylindole) staining in wild type (panels A–D) and *spf2 ctf18* double mutants (panels E–L). Representative spreads are shown for metaphase I (A, E, I), metaphase II (B, F, J, K), anaphase II (C, G), and telophase II (D, H, L). In panel K, a single metaphase II plate is highlighted. White arrowheads indicate meiotic defects, including kinetochore splitting at metaphase I and chromatid splitting, chromatin bridges, lagging chromosomes, or unequal chromatid segregation during meiosis II. The quantification of these defects is presented in Figure S12. Scale bar: 5 µm.

**Figure S12.**
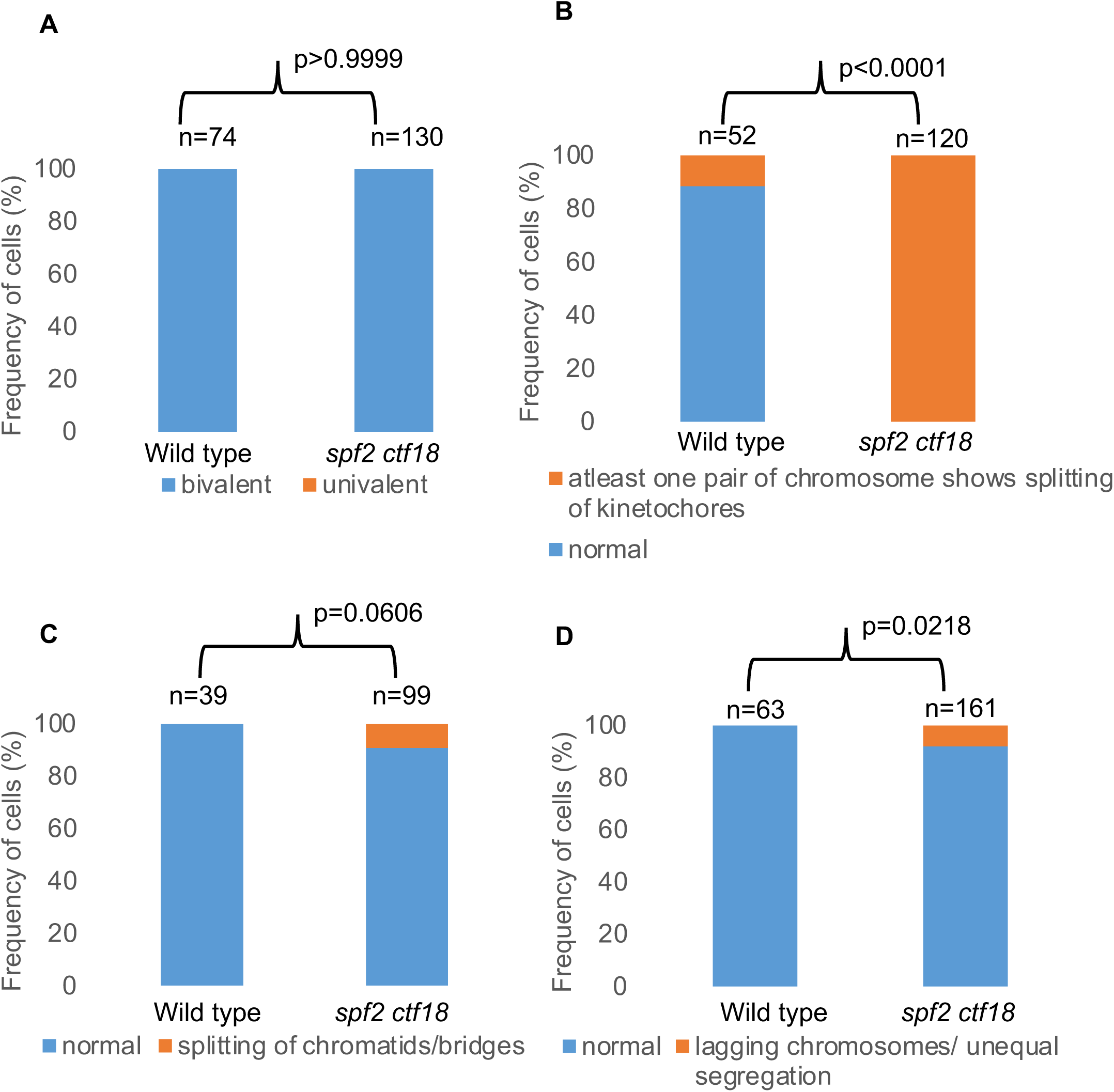
Quantification of meiotic defects in *spf2 ctf18*. (A) Frequency of cells at metaphase I with chromosomes arranged as bivalents (physically connected by chiasmata, indicating crossover formation) or univalents (unpaired chromosomes, suggesting loss of crossover). Bivalents are shown in blue and univalents in orange. The number of cells analyzed is indicated above each bar, and statistical significance was assessed using Fisher’s exact test. (B) Frequency of cells at metaphase I scored for kinetochore shape. Cells without sign of kinetochore splitting are indicated by the blue bar, while cells with at least one chromosome pair showing kinetochore splitting are indicated by the orange bar. (C) Frequency of metaphase II plates, scored for chromatid defects. Blue bar represents normal plates and orange bar indicates cells displaying premature chromatid separation or chromatid bridges. The number of analyzed metaphase II plates is indicated. (D) Frequency of anaphase II- telophase II plates, scored for segregation defects. Blue bar represents normal plates and orange bar indicates plates with unequal chromatid segregation or lagging chromosomes. The Number of analyzed anaphase II plates is indicated

**Figure S13.**
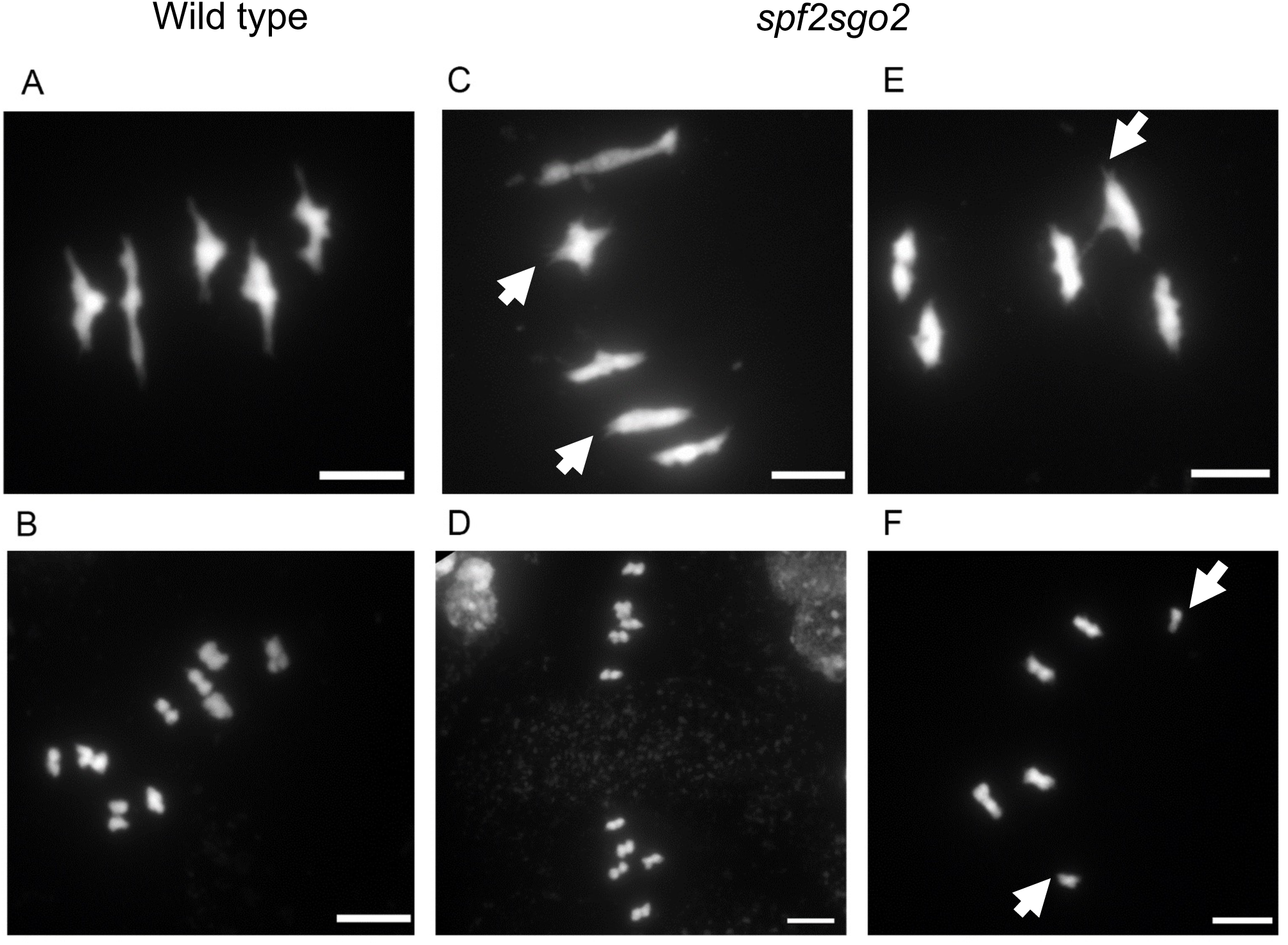
Meiotic Chromosome spreads in wild type and *spf2 sgo2*. Meiotic chromosome spreads were visualized using DAPI (4’, 6-diamidino-2- phenylindole) staining in wild type (panels A – B) and *spf2 ctf18* double mutants (panels C–F). Representative spreads are shown for metaphase I (A, C, E), metaphase II (B, D, F). In panel F, a single metaphase II plate is highlighted. White arrowheads indicate meiotic defects, including kinetochore splitting at metaphase I and chromatid splitting during meiosis II. The quantification of these defects is presented in Figure S14. Scale bar: 5 µm.

**Figure S14.**
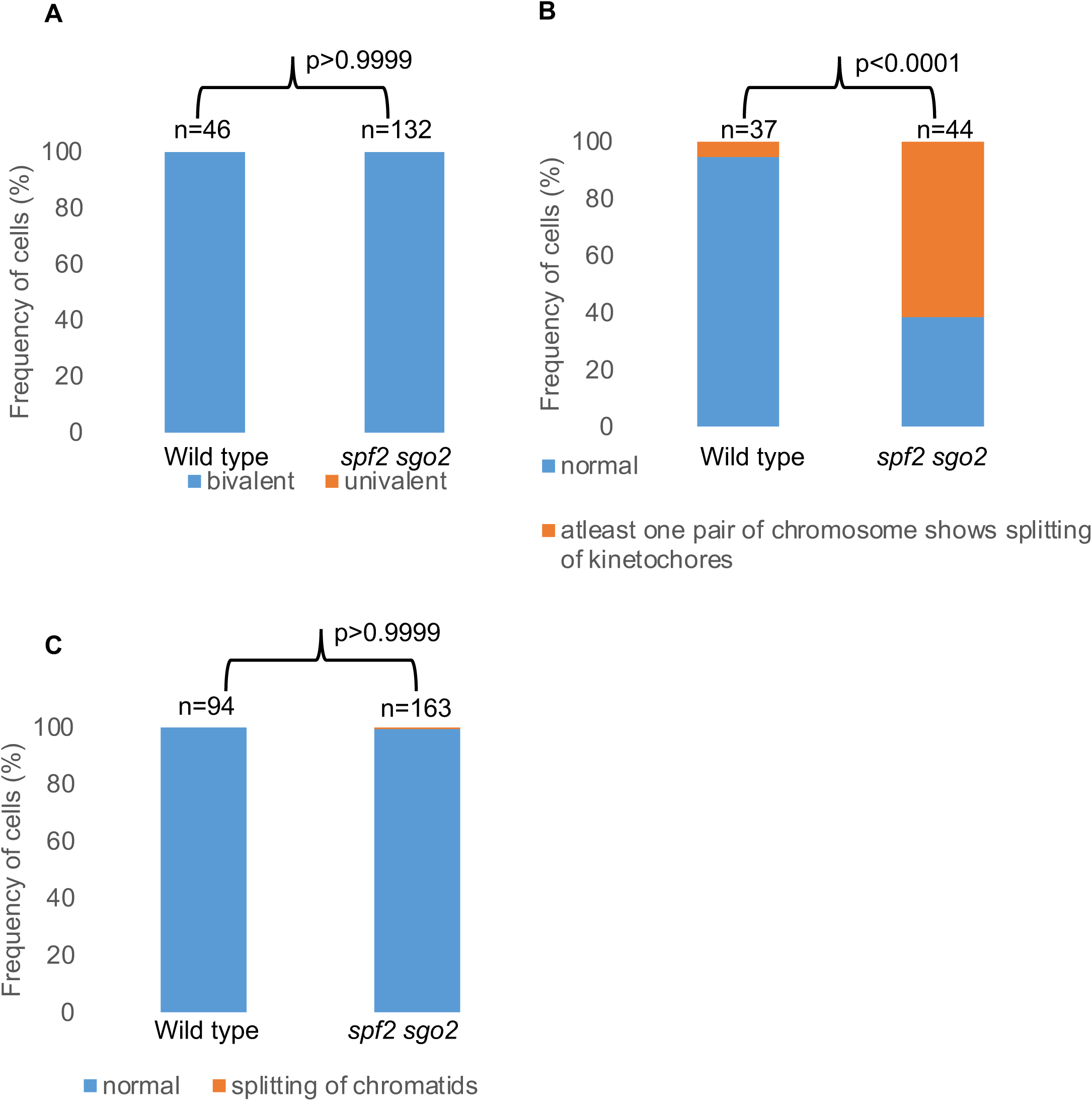
Quantification of meiotic defects in *spf2 sgo2*. (A) Frequency of cells at metaphase I with chromosomes arranged as bivalents (physically connected by chiasmata, indicating crossover formation) or univalents (unpaired chromosomes, suggesting loss of crossover). Bivalents are shown in blue and univalents in orange. The number of cells analyzed is indicated above each bar, and statistical significance was assessed using Fisher’s exact test. (B) Frequency of cells at metaphase I scored for kinetochore shape. Cells without sign of kinetochore splitting are indicated by the blue bar, while cells with at least one chromosome pair showing kinetochore splitting are indicated by the orange bar. (C) Frequency of metaphase II plates, scored for chromatid defects. Blue bar represents normal plates and orange bar indicates cells displaying premature chromatid separation. The number of analyzed metaphase II plates is indicated.

**Supplementary Table S1:** Primers, markers, and guides used in the study.

## Source data

Figure 1A and Figure S2A and S2B – source data – NCBI-Genome assembly ASM2311539v1 (Col-CEN) coordinates of the FTL and PCR markers.

Figure 1B and S2C-G- raw data- recombination frequency measured in cM using CTL3.437.55 interval in single mutants compared to wild type.

Figure 1B and S2C-G- source data- recombination frequency measured in cM using CTL3.437.55 interval in single mutants compared to wild type.

Figure 1C and 1D- raw data- genotyping using PCR markers flanking centromeres to measure recombination frequency in cM in sister wild type of *zyp1, zyp1* mutant, sister wild type of *smc1* and *smc1* mutant.

Figure 1C and 1D- source data-recombination frequency measured in cM using PCR centromere flanking markers in sister wild type of *zyp1, zyp1* mutant, sister wild type of *smc1* and *smc1* mutant.

Figure 2A-source data- Crossover number in various genotypes.

Figure 2B- source data- NCBI-Genome assembly ASM2311539v1 (Col-CEN) coordinates that mark the position of NRZ, LRZ1 and LRZ2.

Figure 2C and Figure 3B- source data- Crossover positions across genome

Figure 3A source data- Crossover frequencies in NRZs and LRZs

Figure 3C. High resolution Crossover position in NRZ/LRZ

Figure 4A and S9-raw data-recombination frequency measured in cM using CTL3.437.55 interval in heterozygous plants obtained from double mutant population.

Figure 4A and S9- source data-recombination frequency measured in cM using CTL3.437.55 interval in heterozygous plants obtained from double mutant population.

Figure 4B and S10-raw data. recombination frequency measured in cM using CTL3.437.55 interval in double mutant population.

Figure 4B and S10- source data-recombination frequency measured in cM using CTL3.437.55 interval in double mutant population.

Figure 5A- raw data- fertility analysis by seed counting in various genotypes.

Figure 5A- source data- fertility analysis by seed counting in various genotypes.

Figure S4- source data- recombination frequency in NRZ, LRZ1 and LRZ2 in wild type and mutants

Figure S12- source data- quantification of meiotic defects in wild type and *spf2 ctf18*

Figure S14- source data- quantification of meiotic defects in wild type and *spf2 sgo2*

## Notes

### Competing Interest Statement

The authors declare the following competing interests: A patent was filed by the Max Planck Society on the use of CTF18, SGO2 and SPF2 to manipulate meiotic recombination in plants, with RM, RSG, JBF, SD, and QL listed as inventors (EP25184806.5. 24.06.2025).

